# Chasing the metabolism of novel syntrophic acetate-oxidizing bacteria in thermophilic methanogenic chemostats

**DOI:** 10.1101/2021.07.06.451242

**Authors:** Yan Zeng, Dan Zheng, Min Gou, Zi-Yuan Xia, Ya-Ting Chen, Masaru Konishi Nobu, Yue-Qin Tang

**Affiliations:** Institute of New Energy and Low-carbon Technology, Sichuan University, No. 24, South Section 1, First Ring Road, Chengdu, Sichuan 610065, China; Biogas Institute of Ministry of Agriculture and Rural Affairs, Section 4-13, Renmin Road South, Chengdu 610041, P. R. China; College of Architecture and Environment, Sichuan University, No. 24, South Section 1, First Ring Road, Chengdu, Sichuan 610065, China; Institute for Disaster Management and Reconstruction, Sichuan University–Hong Kong Polytechnic University, Chengdu, Sichuan 610207, China; Bioproduction Research Institute, National Institute of Advanced Industrial Science and Technology (AIST), Central 6, Higashi 1-1-1, Tsukuba, Ibaraki 305-8566, Japan

**Keywords:** Thermophilic anaerobic digestion, Microbial community, Syntrophic acetate oxidation, Glycine cleavage, Energy conservation

## Abstract

**Background:** Acetate is the major intermediate of anaerobic digestion of organic waste to CH_4_. In anaerobic methanogenic systems, acetate degradation is carried out by either acetoclastic methanogenesis or a syntrophic degradation by a syntrophy of acetate oxidizers and hydrogenotrophic methanogens. Due to challenges in isolation of syntrophic acetate-oxidizing bacteria (SAOB), the diversity and metabolism of SAOB, as well as the mechanisms of their interactions with methanogenic partners remain poorly understood.

**Results:** In this study, we successfully enriched previously unknown SAOB by operating continuous thermophilic anaerobic chemostats fed with acetate, propionate, butyrate, or isovalerate as the sole carbon and energy source. They represent novel clades belonging to Clostridia, Thermoanaerobacteraceae, Anaerolineae, and Gemmatimonadetes. In these SAOB, acetate is degraded through reverse Wood-Ljungdahl pathway or an alternative pathway mediated by the glycine cleavage system, while the SAOB possessing the latter pathway dominated the bacterial community. Moreover, H_2_ is the major product of the acetate degradation by these SAOB, which is mediated by [FeFe]-type electron-confurcating hydrogenases, formate dehydrogenases, and NADPH reoxidation complexes. We also identified the methanogen partner of these SAOB in acetate-fed chemostat, *Methanosarcina thermophila*, which highly expressed genes for CO_2_-reducing methanogenesis and hydrogenases to supportively consuming H_2_ at transcriptional level. Finally, our bioinformatical analyses further suggested that these previously unknown syntrophic lineages were prevalent and might play critical roles in thermophilic methanogenic reactors.

**Conclusion:** This study expands our understanding on the phylogenetic diversity and *in situ* biological functions of uncultured syntrophic acetate degraders, and presents novel insights on how they interact with their methanogens partner. These knowledges strengthen our awareness on the important role of SAO in thermophilic methanogenesis and may be applied to manage microbial community to improve the performance and efficiency of anaerobic digestion.

## Background

Anaerobic digestion (AD) of organic waste to produce methane offers opportunities to deliver multiple environmental benefits as it encompasses organic waste treatment and renewable energy production. Volatile fatty acids (VFAs) are the main intermediates, and thus syntrophic fatty acid oxidation is thought to be the key step in AD [1]. Notably, acetate serves as the most important intermediate metabolite and the major precursor of methane, accounting for 60 to 80% of methane production in anaerobic digesters [2, 3]. Notably, metabolic disorders would lead to accumulation of acetate in anaerobic digesters, which may cause acidification and reduce methane production, destabilizing the AD systems. Therefore, uncovering the underly mechanism of anaerobic acetate metabolism is fundamental to manage microbial AD system for better performance.

Under methanogenic condition, methane production from acetate follows two routes (i) acetoclastic methanogenesis (Eq. 1), (ii) syntrophic acetate oxidation coupled with hydrogenotrophic methanogenesis (Eqs. 2 and 3) [4].

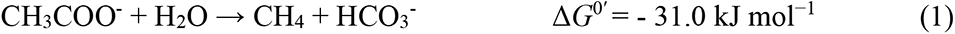

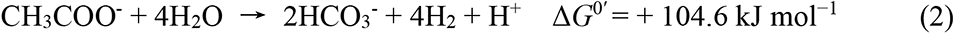

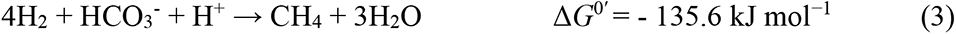

In the former route, acetate is cleaved into carbonyl group and methyl group, then respectively oxidized to CO_2_ and reduced to CH_4_ by aceticlastic methanogens *Methanosarcina* or *Methanothrix* [5]. In the latter route, both methyl and carbonyl group of acetate are oxidized to CO_2_, associated with the generation of H_2_. This reaction is thermodynamically unfavorable under standard conditions (ΔG^0’^= +104.6 kJ mol^-1^). Thus, “syntrophic” cooperation with hydrogen-scavenging methanogenic partners (ΔG^0’^= -135.6 kJ mol^-1^) is necessary to maintain thermodynamic favorability [4, 6]. Previous studies report observation of syntrophic acetate oxidation in selective conditions (*e.g.*, high concentration of ammonia [7], high temperature [8], or low loading rate and long retention time [9]), suggesting that this niche may play an critical role in diverse methanogenic systems that may have challenges in supporting aceticlastic methanogens.

Although six strains of syntrophic acetate-oxidizing bacteria (SAOB) have been cultured, the full diversity and metabolism of SAOB still remain poorly understood. Among described SAOB, while three species (*Thermacetogenium phaeum* [10], *Syntrophaceticus schinkii* [11], and *Tepidanaerobacter acetatoxydans* [12]) possess the well-known reverse Wood-Ljungdahl (WL) pathway for syntrophic acetate oxidation [13, 14], but two (*Pseudothermotoga lettingae* [15] and *Schunerera ultunensis,* previously *Clostridium ultunense* [16]) lack genes for this classical WL pathway and are suspected to possess an alternative metabolism, potentially mediated by a glycine cleavage system pathway [13, 17]. Moreover, these cultured SAOB are generally detected with low abundances in anaerobic bioreactors [18–21]. Previous studies based on DNA or protein stable isotope probing (SIP) point towards the presence of other phylogenetically distinct uncultured acetate oxidizers in anaerobic digestors [22–24]. Therefore, uncovering the diversity, ecology, metabolism, and symbiotic interactions of these yet-to-be cultured SAOB is essential for improving our understanding and operation of methanogenic bioreactors under stressed conditions.

Furthermore, it is also crucial for understanding the mechanisms of the syntrophic interaction between SAOB and their methanogens partner. Interspecies electron transfer between these two groups is essential to maintain thermodynamic favorability of microbial methanogenesis [25, 26]. Previous survey on syntrophic acetate metabolizers suggested that H_2_ has been regarded as electron carrier from acetate oxidizers to methanogens [4, 14, 27]. Several recent studies also suggested that formate transfer also play a role in propionate [28, 29] and isovalerate [30] syntrophic degradation. Direct interspecies electron transfer (DIET) activity has been suggested in enrichment communities degrading propionate and butyrate [31, 32]. However, whether formate transfer and DIET transfer play roles in acetate syntrophic degradation is not yet known.

One effective cultivation-independent approach to studying the physiology and *in situ* metabolism of uncultured organisms is the combination of metagenomics and metatranscriptomics [17, 30, 33, 34]. In this study, we employed such approach to recover genomes (metagenome-assembled genomes, MAGs) and gene expression profiles of novel potential acetate degraders from fatty-acid-fed thermophilic anaerobic chemostats to investigate their catabolic pathways, energy conservation, and metabolic interactions with their methanogens partner.

## Results and discussions

### Chemostat operation and performance

Eight thermophilic anaerobic chemostats were stably operated with synthetic wastewater containing acetate, propionate, butyrate, or isovalerate as the sole carbon and energy source at different loading rates (Table S1; Materials and Methods). During the steady operation period, the performance of chemostats was stable, *i.e*., biogas production was stable and concentrations of VFAs in the eight chemostats were markedly low (10∼30 mg L^-1^), indicating that VFAs fed were almost completely degraded by these microbial communities (Table 1 and Fig. S1).

**Table 1.** Operational performance of mesophilic and thermophilic chemostats during the steady operation period ^a^.

| Chemostat name | ATL | ATH | PTL | PTH | BTL | BTH | VTL | VTH |
| --- | --- | --- | --- | --- | --- | --- | --- | --- |
| Carbon source | Acetate | Acetate | Propionate | Propionate | Butyrate | Butyrate | Isovalerate | Isovalerate |
| Dilution rate (day<sup>-1</sup>) | 0.025 | 0.05 | 0.025 | 0.05 | 0.025 | 0.05 | 0.025 | 0.05 |
| HRT (day) | 40 | 20 | 40 | 20 | 40 | 20 | 40 | 20 |
| Inflow concentration (mg·L<sup>-1</sup>) | 20000 | 20000 | 16444 | 16444 | 14667 | 14667 | 13600 | 13600 |
| Gas production rate (mL·L<sup>-1</sup>·d<sup>-1</sup>) | 115±22 | 294±67 | 103±17 | 383±43 | 155±28 | 396±35 | 143±37 | 365±18 |
| CH<sub>4</sub> content (%) | 58±4 | 60±5 | 62±2 | 63±3 | 66±2 | 73±4 | 67±4 | 76±6 |
| H<sub>2</sub> partial pressure (Pa) | 0.8±0.4 | 1.4±0.6 | 2.9±1.2 | 3.1±1.3 | 2.3±1.4 | 2.9±1.3 | 2.0±1.2 | 2.5±0.9 |
| pH | 8.04±0.14 | 7.85±0.14 | 7.95±0.14 | 7.78±0.14 | 7.78±0.08 | 7.63±0.11 | 7.84±0.14 | 7.66±0.10 |
| TOC (mg·L<sup>-1</sup>) | 139±83 | 132±78 | 84±49 | 118±46 | 87±74 | 123±91 | 84±49 | 114±44 |
| Formate (mg·L<sup>-1</sup>)<sup>b</sup> | ND | ND | ND | ND | ND | ND | ND | ND |
| Acetate (mg·L<sup>-1</sup>) | 11±22 | 12±35 | 6±8 | 2±4 | 31±47 | 27±69 | 23±53 | 20±20 |
| Propionate (mg·L<sup>-1</sup>) | 0 | 0 | 11±13 | 15±48 | ~1.0 | ~1.0 | ~1.0 | ~1.0 |
| Butyrate (mg·L<sup>-1</sup>) | 0 | 0 | 0 | 0 | ~1.0 | ~1.0 | ~1.0 | ~1.0 |
| Valerate (mg·L<sup>-1</sup>) | 0 | 0 | 0 | 0 | 0 | 0 | ~1.0 | ~1.0 |
| Iso-valerate (mg·L<sup>-1</sup>) | 0 | 0 | 0 | 0 | 0 | 0 | ~1.0 | ~1.0 |
| SS (g·L<sup>-1</sup>) | 0.421±0.08 | 0.543±0.14 | 0.468±0.11 | 0.68±0.07 | 1.43±0.49 | 1.44±0.38 | 1.35±0.44 | 1.38±0.25 |
| VSS (g·L<sup>-1</sup>) | 0.308±0.07 | 0.392±0.11 | 0.352±0.10 | 0.513±0.07 | 0.62±0.21 | 0.65±0.18 | 0.57±0.21 | 0.67±0.19 |

<sup>a</sup> HRT, hydraulic retention time; TOC, total organic carbon; VSS, volatile suspended solid; ND, not detected. The operational parameters (*e.g.*, pH, gas production rate, and CH<sub>4</sub> content, etc.) were the averages (mean ± SD, n > 30) during the steady-state period in ATL (day 100–550), ATH (day 80–550), PTL (day 100– 650), PTH (day 300–450), BTL (day 100–650), BTH (day 300–650), VTL (day 150–650), AND ATL (day 300–650) chemostats (Fig. S2). <sup>b</sup> Formate concentration was below the detective limit (10 mg L<sup>-1</sup>) of the high-performance liquid chromatography.

### Microbial diversity and community composition of thermophilic anaerobic chemostats

Based on DNA- and RNA-based 16S ribosomal RNA gene analysis, the bacterial community of the thermophilic chemostats contain diverse population belonging to uncultured lineages (Figs. 1 and S2). The dominated bacterial populations included *Firmicutes* (*e.g.*, order MBA03 and family Thermoanaerobacteraceae), *Bacteroidetes* (Lentimicrobiaceae), and *Chloroflexi* (Anaerolineaceae), which were at high abundance (up to 72%) and activity (up to 41%) in the all the thermophilic chemostats. *Thermodesulfovibrio* displayed low DNA-based relative abundance, but also high activity (up to 18% of transcriptome). Notably, one genus associated with previously isolated syntrophic acetate-oxidizing bacteria (SAOB) (*Tepidanaerobacter*; [12]) was detected but only comprised less than 1% of the total bacterial community.

**Fig. 1.**
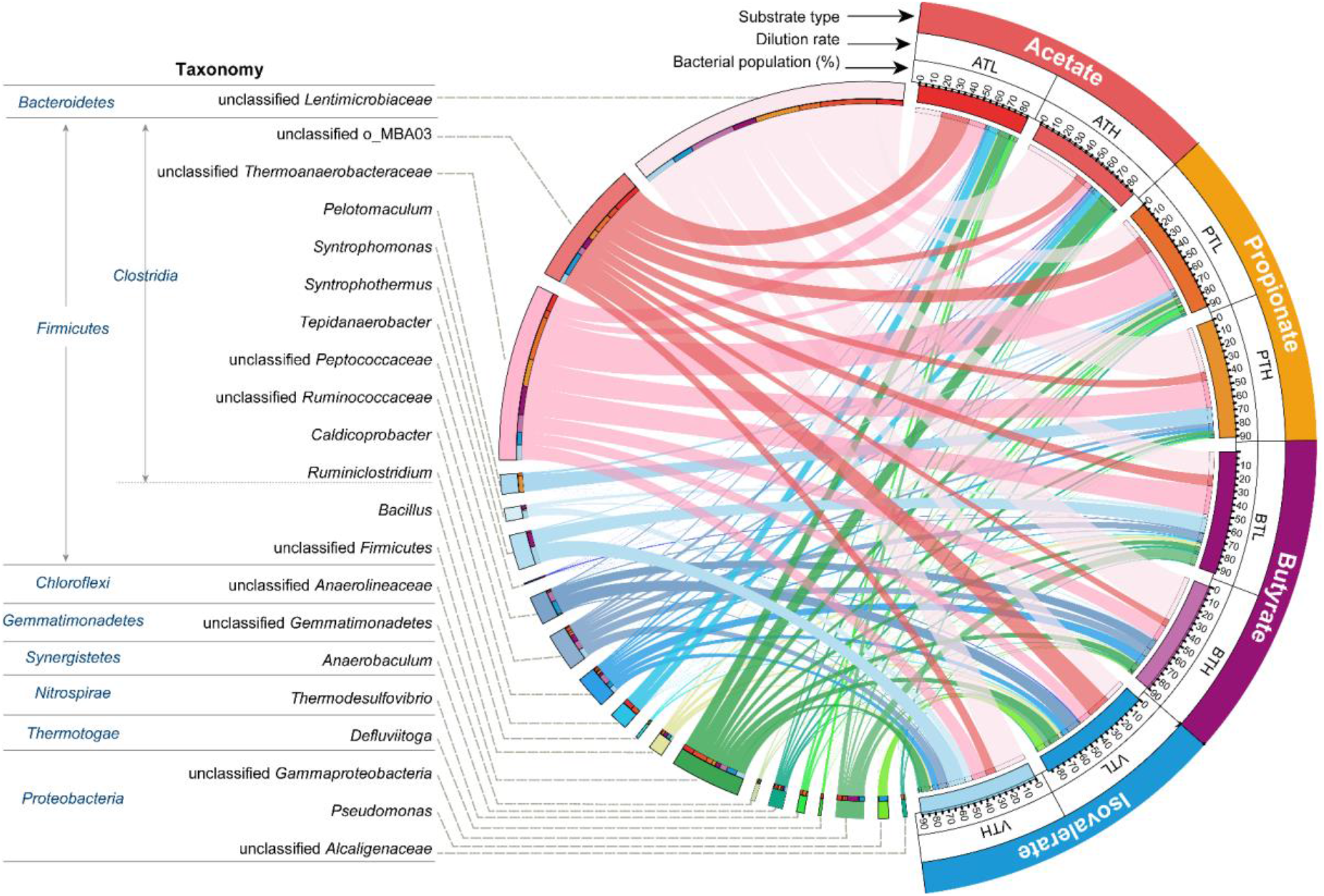
Relative abundance of bacterial genera based on 16S rRNA gene amplicon sequencing in the thermophilic chemostats (DNA level).

In regarding to archaeal community, according to 16S rRNA gene analysis, *Methanosarcina* whose relative abundance accounted for 26%∼94% and 48%∼99% of archaeal community at the DNA and RNA level, respectively, was the active methanogen across all the thermophilic chemostats. *Methanosarcina* OTUs held a 99.5% sequence similarity to multitrophic methanogen *Methanosarcina thermophila* TM-1, which is able to turnout H_2_/formate, acetate, methanol to methane [35]. *Methanothermobacter* was dominant in PTL (67%-DNA, 51%-RNA), PTH (47%-DNA, 42%-RNA), and BTL (25%-DNA, 24%-RNA), while *Methanoculleus* predominated in BTH (33%-DNA, 29%-RNA) and VTH (59%-DNA, 9%-RNA) (Fig. S4).

To profile the metabolic capability of such bacteria and methanogens (potential partners and competitors), a total of 173 Gbp metagenomic clean sequences (ATL, 35 Gbp; PTL, 69 Gbp; BTL, 34 Gbp; VTL, 35 Gbp) were obtained. Illumina paired-end reads from the two subsamples (three subsamples in PTL) were co-assembled. Binning the assembled contigs of metagenomes for thermophilic communities yielded 108, 157, 81 and 96 MAGs from ATL, PTL, BTL, and VTL, respectively. To obtain gene expression profiles of the bacteria and archaea in the chemostats, a total of 287 million metatranscriptomic reads (33.6 Gbp, approximately 4.2 Gbp for each RNA sample) were sequenced and mapped to the MAGs for each chemostat (78–95% of reads mapped using a 100% nucleotide similarity cutoff). Based on mapping metagenomic reads to the obtained MAGs, the bacterial populations retrieved accounted for 79%, 52%, 67% and 67% of the metagenomic reads obtained from ATL, PTL, BTL and VTL (Fig. S3C). In addition, these bacterial populations accounted for 80%, 74%, 65% and 73% of the metatranscriptomic reads from ATL, PTL, BTL and VTL, respectively (Fig. S3D).

As for the methanogen archaea, *Methanosarcina thermophila* (MAG.ATL014) was the active methanogen in ATL, PTL, BTL and VTL, accounting for 18%, 11%, 3% and 15% of the metatranscriptomic reads from PTL, BTL and VTL, respectively (Fig. S5 and Table S2). The activity of hydrogenotrophic methanogens was negligible in ATL. *Methanothermobacter* (MAG.PTL002) was dominant accounting for 13% of the metatranscriptomic reads in PTL. *Methanoculleus* (MAG.BTL076 and MAG.VTL077) predominated in BTL and VTL, accounting for 28% and 8% of the metatranscriptomic reads from BTL and VTL, respectively.

### Syntrophic metabolism and energy conservation of acetate-degrading community in ATL

In the analysis of community structure based on 16S rRNA gene sequencing, the relative abundance and RNA-based activity of bacteria was greater than that of archaea in the all the eight chemostats (Fig. S3A and S3B). This phenomenon was also observed in the analysis based on metagenome and metatranscriptome data (Fig. S3C and S3D). In methanogenic system, syntrophic fatty acid oxidizers convert propionate, butyrate, and isovalerate to acetate and H_2_/formate, and symbiotically hand off these by-products to partnering acetate- and H_2_-consuming methanogenic archaea [23, 28, 30]. Therefore, it was expected that bacteria displayed high abundance and activity in propionate-, butyrate-, and isovalerate-fed chemostats, which is consistent with our observation. In acetate-fed chemostats, since aceticlastic methanogens, such as *Methanosarcina*, could autonomously degraded acetate, they were expected to dominate the methanogenic communities. However, despite that *Methanosarcina* (99.5% rRNA sequence similarity to *M. thermophila* TM-1; MAG: MAG.ATL014) held a considerably high abundance and activity in our acetate-fed chemostats, the community are unexpectedly dominated by bacterial populations. This result suggested that several bacterial populations may play a significant role in acetate degradation in our acetate-fed chemostats, and potentially be the previously unknown SAOB clades that we were looking for.

#### High activity of CO_2_-reducing methanogenesis in Methanosarcina

Although *Methanosarcina* groups were previously reported as the main competitor of SAOB, since bacterial populations largely dominated the whole community, we hypothesized that *Methanosarcina* groups could also utilize bacteria-produced metabolites (*e.g.*, H_2_ and CO_2_) to produce methane in our acetate-fed chemostats, playing a role as the partner of the potential SAOB in the community. To test this hypothesis, we first analyzed the metabolic feature of the *Methanosarcina* MAGs in ATL. In accordance with our hypothesis, the dominant *Methanosarcina* MAG.ATL014 interestingly expressed genes for CO_2_ reduction (in addition to acetate catabolism; top octile and quartile of expressed genes in the corresponding MAG respectively; Figs. 2 and 3B; Table S3) even though *Methanosarcina* sp. are known to significantly downregulate expression of such genes during acetate degradation as they are not necessary [36, 37]. Decrease in the activity of the CO_2_-reducing pathway also results in decreased cellular concentrations (up to 10-fold) of coenzyme F_420_, an electron carrier for the CO_2_ branch during growth on acetate [38], and, though qualitative, *Methanosarcina*-like cells showed higher autofluorescence (at 420 nm) in chemostats where *Methanosarcina* highly expressed the CO_2_-reducing pathway (*i.e.*, ATL and ATH compared to PTL in Fig. S6; [28]). Thus, the *Methanosarcina in situ* likely utilizes an alternative electron source in parallel with acetate.

**Fig. 2.**
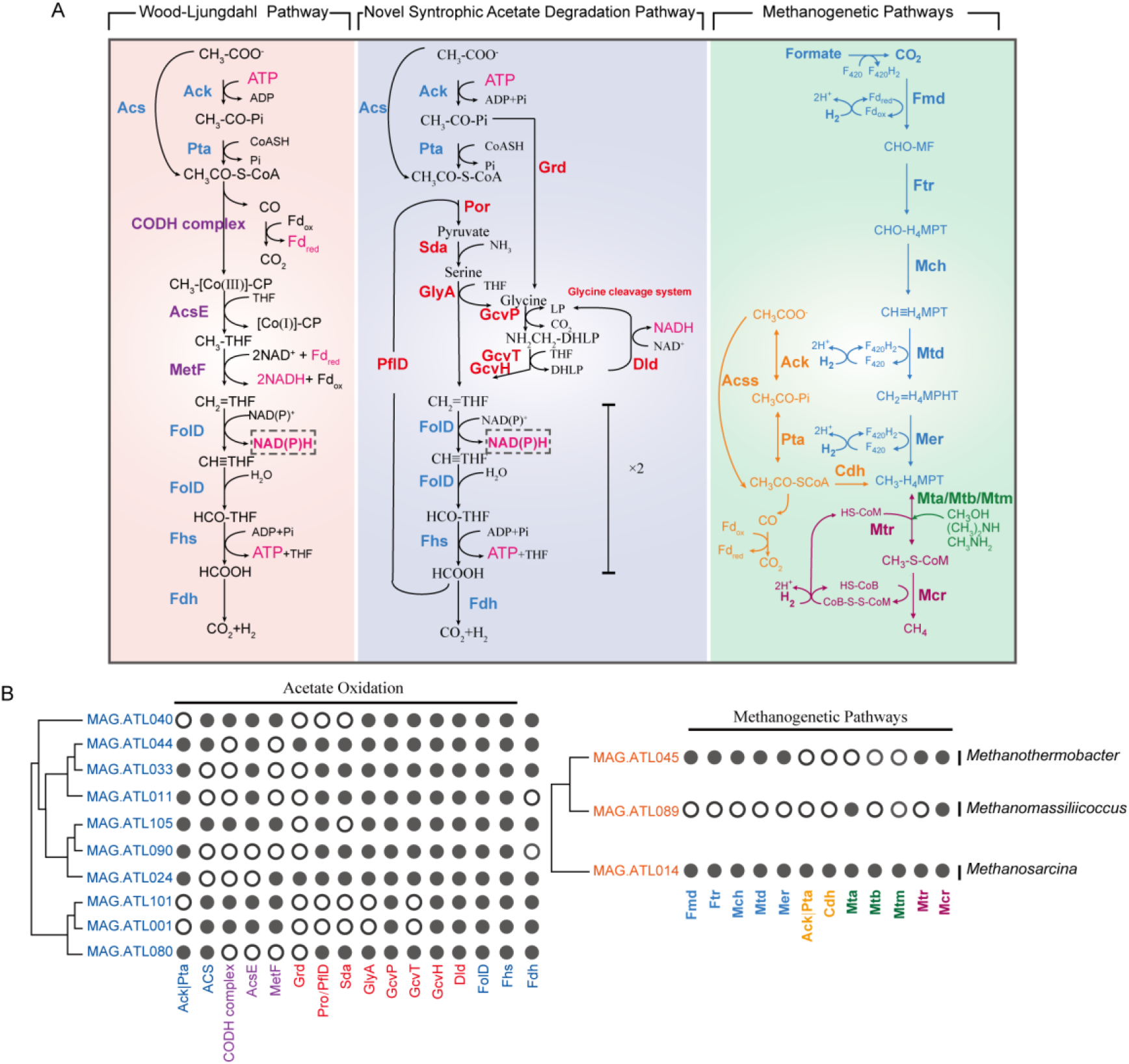
Metabolic pathways of acetate oxidation and methanogenesis in thermophilic acetate-degrading chemostat (A), and distribution of catabolic pathways among the studied contributors (B). For each syntroph and methanogen, we show presence (indicated by filled circle) of genes encoding pathways for acetate catabolism and methanogenesis. Enzyme abbreviations are as follows: Ack, acetate kinase; Pta, phosphate acetyltransferase; Acs, acetyl-CoA synthetase; CODH complex, acetyl CoA synthetase complex; AcsE, methyltetrahydrofolate:corrinoid/iron-sulfur protein methyltransferase; MetF, methylenetetrahydrofolate reductase; FolD, methylenetetrahydrofolate dehydrogenase/cyclohydrolase; Fhs, formate--tetrahydrofolate ligase; Fdh, formate dehydrogenase. Grd, glycine reductase; Por, pyruvate dehydrogenase; PflD, pyruvate-formate lyase; Sda, serine dehydratase; GlyA, glycine hydroxymethyltransferase; GcvPA, glycine dehydrogenase subunit A; GcvPB, glycine dehydrogenase subunit B, GcvT, glycine cleavage system T protein; GcvH, glycine cleavage system H protein; Dld, dihydrolipoyl dehydrogenase. Fmd, formylmethanofuran dehydrogenase; Ftr, formylmethanofuran-tetrahydromethanopterin N-formyltransferase; Mch, methenyltetrahydromethanopterin cyclohydrolase; Mtd, methylenetetrahydromethanopterin dehydrogenase; Mer, F_420_-dependent 5,10-methenyltetrahydromethanopterisn reductase; Cdh, acetyl-CoA decarbonylase/synthase complex; Mta, [methyl-Co(III) methanol-specificcorrinoid protein]:CoM methyltransferase; Mtb, [methyl-Co(III) dimethylamine-specificcorrinoid protein]:CoM methyltransferase; Mtm, [methyl-Co(III) monomethylamine-specificcorrinoid protein]:CoM methyltransferase ; Mtr, tetrahydromethanopterin S-methyltransferase; Mcr, methyl-CoM reductase. CHO-MF, formyl-methanofuran; CHO-H_4_MPT, formyl-tetrahydromethanopterin; CH≡H_4_MPT, methenyl-tetrahydromethanopterin; CH_2_=H_4_MPT, methylene-tetrahydromethanopterin; CH_3_-H_4_MPT, methyl-tetrahydromethanopterin; CH_3_-S-CoM, methyl-coenzyme M; HS-CoM, coenzyme M; HS-CoB, coenzyme B; CoB-S-S-CoM, mixed disulphide of CoM and CoB. Enzyme abbreviations and their corresponding genes are elaborated in Supporting Information Tables S3 and S5.

**Fig. 3.**
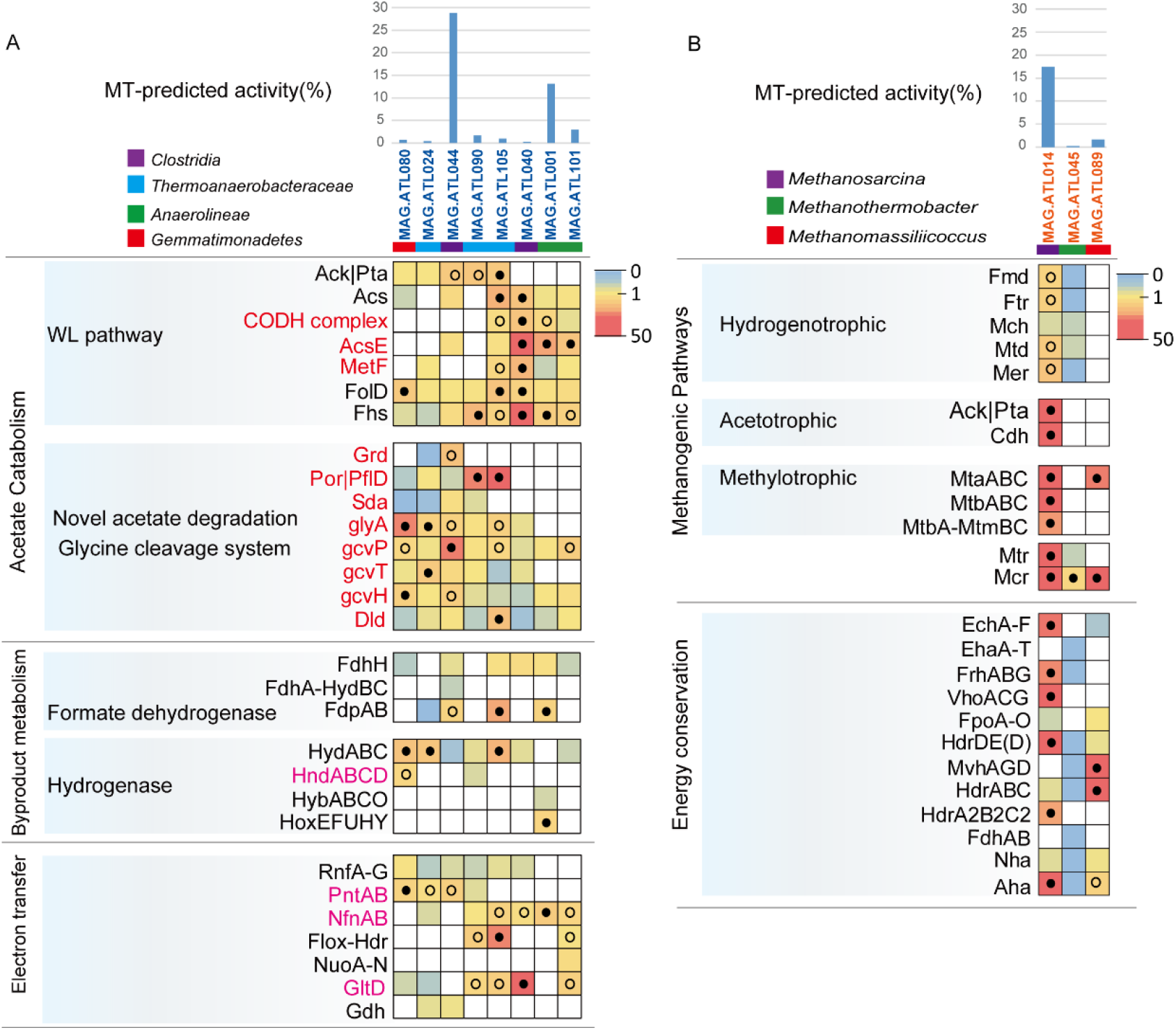
Gene expression level for acetate oxidation, H_2_/formate metabolism, and electron transfer genes in syntrophs which may syntrophically degrade acetate (A) and methanogens (B) from thermophilic acetate-degrading chemostat. For each metagenome-assembled genome (MAG), the percentages of the metatranscriptomic (MT) reads mapped to the MAG out of the metatranscriptomics mapped to all MAGs (both *Bacteria* and *Archaea*) are shown. The gene expression levels are calculated as reads per kilobase of transcript per million reads mapped to individual MAG (RPKM) normalized to the median gene expression for the corresponding MAG (RPKM-NM) averaged from duplicate samples. Pathways containing genes with RPKM-NM greater than the octile and quartile are marked (filled and open dots, respectively). Enzyme abbreviations and their corresponding genes are elaborated in Supporting Information Tables S3-S6.

Given the lack of exogenous H_2_ and methylated compounds (*i.e.*, compounds that would stimulate usage of the CO_2_-reducing branch), this may indicate the presence of some bacterial populations in the chemostat catabolizing acetate and syntrophically transferring H_2_ and/or electrons to *Methanosarcina*. Supporting this, (i) acetate-degrading *M. thermophila* cells are known to consume H_2_ with affinity similar to that of hydrogenotrophic methanogens [39] and (ii) *Methanosarcina* MAG.ATL014 highly expressed hydrogenases (Fig. 3B; Table S4). This is consistent with previous studies that *Methanosarcina* had been observed together with SAOB in acetate-fed thermophilic anaerobic digesters [22, 40]. Moreover, the detection of H_2_-utilizing methanogens (*Methanothermobacter* MAG.ATL045 and *Methanomassiliicoccus* MAG.ATL089; albeit at much lower activity levels) also suggests that H_2_ transfer is taking place *in situ* (Figs. 2 and 3B; Table S3). The parallel expression of methanol-reducing methanogenesis by *Methanomassiliicoccus* and methyl compound metabolism by *Methanosarcina* suggests that *Methanosarcina* may generate methanol (methanol:coenzyme M methytransferase has been shown to generate methanol *in vitro barkeri* [41]) and feed into *Methanomassiliicoccus* methanogenesis.

#### Putative syntrophic acetate metabolizers

To identify potential uncultured SAOB that may interact with the above methanogens, we performed metabolic reconstruction of the MAGs recovered for abundant and active bacterial populations. Genome- and transcriptome-based prediction of SAOB is challenging given that the conventional acetate oxidation pathway (reverse WL pathway) and the previously proposed glycine-mediated alternative pathway can be used for carbon fixation and serine/glycine biosynthesis respectively. To identify genotypic features associated with SAOB, we performed comparative genomics of isolated SAOB. All isolated SAOB that possess the reverse WL pathway conserve NAD(P) transhydrogenase, while organisms only capable of homoacetogenesis do not encode genes for this enzyme. Both SAOB (*S. ultunensis* and *P. lettingae*) that lack the WL pathway possess NADPH re-oxidizing complexes, albeit different enzymes: NADPH-dependent FeFe hydrogenase (*S. ultunensis*) and NADH-dependent NADP:ferredoxin oxidoreductase (*P. lettingae*). *P. lettingae* has previously been proposed to use a glycine dehydrogenase-mediated pathway for C1 metabolism and *S. ultunensis* may also use this pathway as it lacks the conventional reverse WL pathway. Interestingly, *S. ultunensis* encodes the glycine dehydrogenase directly upstream of NADPH-dependent FeFe hydrogenase, suggesting potential association of glycine metabolism, NADPH reoxidation, and H_2_ generation. Thus, we restricted our analysis to populations encoding and highly expressing the WL pathway or glycine-mediated pathway along with NADPH re-oxidation and H_2_/formate generation (expression in top quartile of each population’s expression profile). To further increase the stringency of our analysis, we further exclude any populations that highly express amino acid catabolism (which is often NADP-dependent) using expression of glutamate dehydrogenase as a marker (i.e., gdhA/gdhB in top quartile of expression profile).

We identified bacterial populations associated with uncultured Clostridia (MAG.ATL040, MAG.ATL011, MAG.ATL033, and MAG.ATL044), Thermoanaerobacteraceae (MAG.ATL024, MAG.ATL090, and MAG.ATL105), Anaerolineae (MAG.ATL001 and MAG.ATL101) and Gemmatimonadetes (MAG.ATL080) as the potential SAOB that encode the reverse WL and glycine-mediated acetate-oxidizing pathways and complementary NADPH re-oxidation and H_2_/formate-generating enzymes (Figs. 2 and S7; Tables S5 and S6) [17, 42]. Phylogenetic analysis revealed that these bacterial populations were distantly related to known SAOB (Fig.4). Furthermore, the Clostridia members (MAG.ATL011, MAG.ATL033, and MAG.ATL044) and Thermoanaerobacteraceae members (MAG.ATL024, MAG.ATL090, and MAG.ATL105) were phylogenetically closely related to each other, but distantly related to any cultured organisms (Fig.4). Based on the above criteria, among these populations, Thermoanaerobacteraceae population (MAG.ATL105), Anaerolineae populations (MAG.ATL001 and MAG.ATL101) and *Clostridia* population (MAG.ATL040) may syntrophically degrade acetate via reverse Wood-Ljungdahl pathway. Thermoanaerobacteraceae population (MAG.ATL024 and MAG.ATL090), *Clostridia* population (MAG.ATL044) and Gemmatimonadetes (MAG.ATL080) may syntrophically degrade acetate via *Thermotogae*-associated pathway; Figs. 2 and 3A; Tables S5 and S6). These results suggested that previously unknown bacterial clades plays a critical role in syntrophic acetate oxidation.

**Fig. 4.**
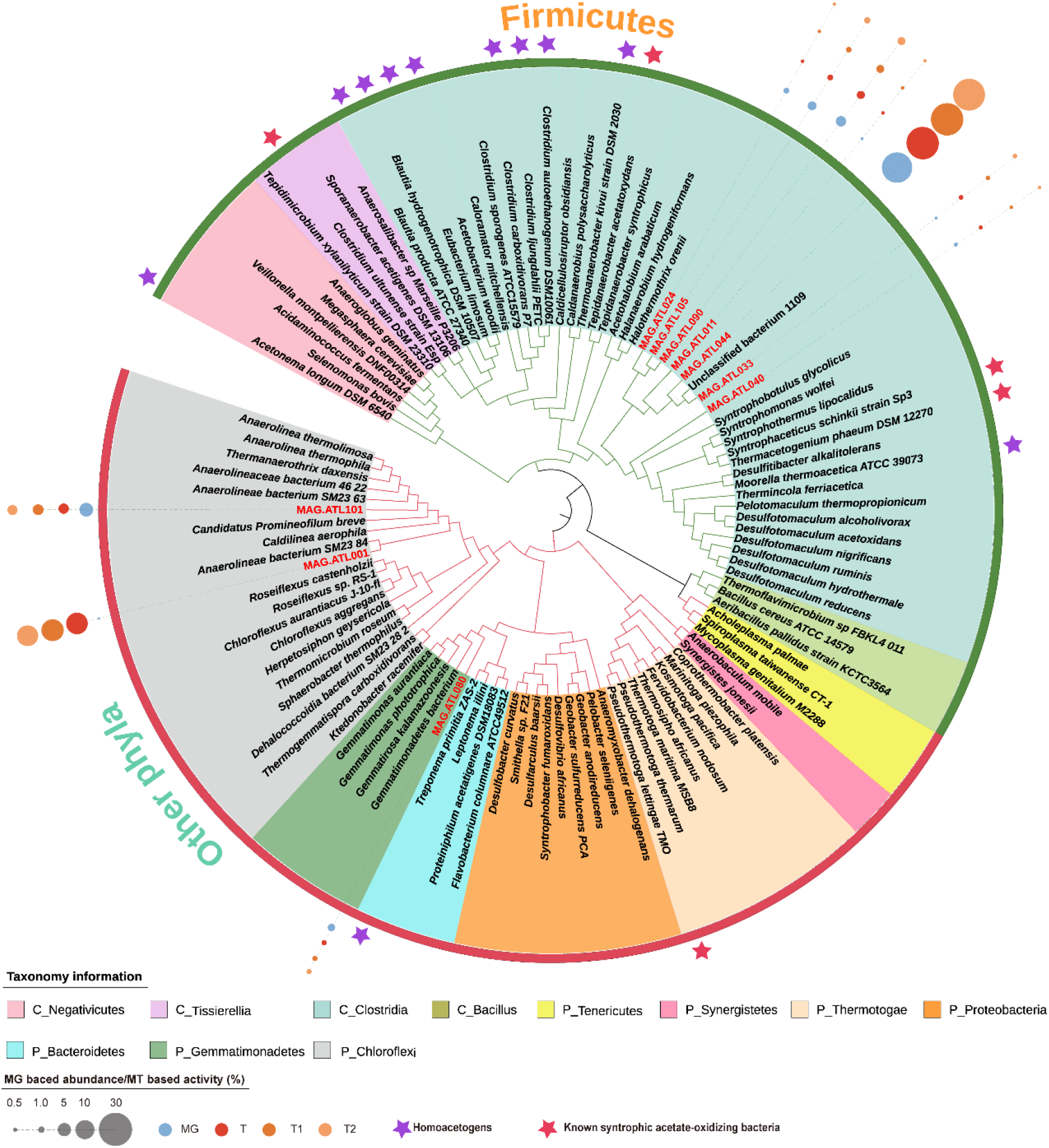
Phylogenetic analyses of metagenome-assembled genomes (MAGs) of syntrophs in thermophilic acetate-degrading chemostat. The corresponding abundance of MAGs are estimated from their metagenomic coverage calculated as the percentage of metagenomics (MG) reads mapped to each MAG relative to the total reads mapped to all bacterial and archaeal MAGs. The estimated activity of MAGs in acetate-degrading chemostat are shown as the percentage of metatranscriptomic (MT) reads mapped to each MAG relative to total reads mapped to all bacterial and archaeal MAGs (T, totoal MT reads; T1, MT reads of sampling time point 1; T2, MT reads of sampling time point 2).

#### Energy conservation and electron flow in acetate oxidizers

Metabolism under methanogenic conditions necessitates complementation of substrate oxidation with electron balance. Thus, we explored energy conservation systems (*e.g.*, electron transfer and electron confurcation/bifurcation) of the putative acetate oxidizers. Most of the putative SAOB encode cytoplasmic [FeFe]-type electron-confurcating hydrogenases (HydABC) (Figs. 3A and 5; Table S6) that use exergonic oxidation of reduced ferredoxin (Fd_red_) (E^0’^ = −430 mV) to drive unfavorable H_2_ generation from NADH oxidation (E^0’^ = −230 mV) [43], a energy conservation strategy associated with syntrophic fatty acid oxidizers [14, 29, 44]. In addition, populations (MAG.ATL080 and MAG.ATL090) performed H_2_ generation via the NADPH-dependent [FeFe] hydrogenase (HndABCD) (Figs. 3A and 5; Table S6). The Anaerolineae member (MAG.ATL001) also encodes a cytochrome b-linked NiFe hydrogenase (HybABCO) and a cytosolic NiFe hydrogenase (HoxEFUHY) (Figs. 3A and 5; Table S6). As for formate metabolism, six of eight SAOB MAGs possess a ferredoxin-dependent formate dehydrogenase (FdhH). The Clostridia-related member MAG.ATL044, Thermoanaerobacteraceae (MAG.ATL105), and Anaerolineae (MAG.ATL001) highly expressed a putative NADPH-dependent formate dehydrogenases (FdpAB). MAG.ATL044 harbors a putative NAD^+^-dependent electron-bifurcating complex (FdhA-hydBC: formate dehydrogenase organized with HydBC-related subunits) (Figs. 3A and 5; Table S6). Therefore, the putative SAOB identified encode and express enzymes for energy conservation and electron flow that support thermodynamically challenging catabolism and syntrophy.

**Fig. 5.**
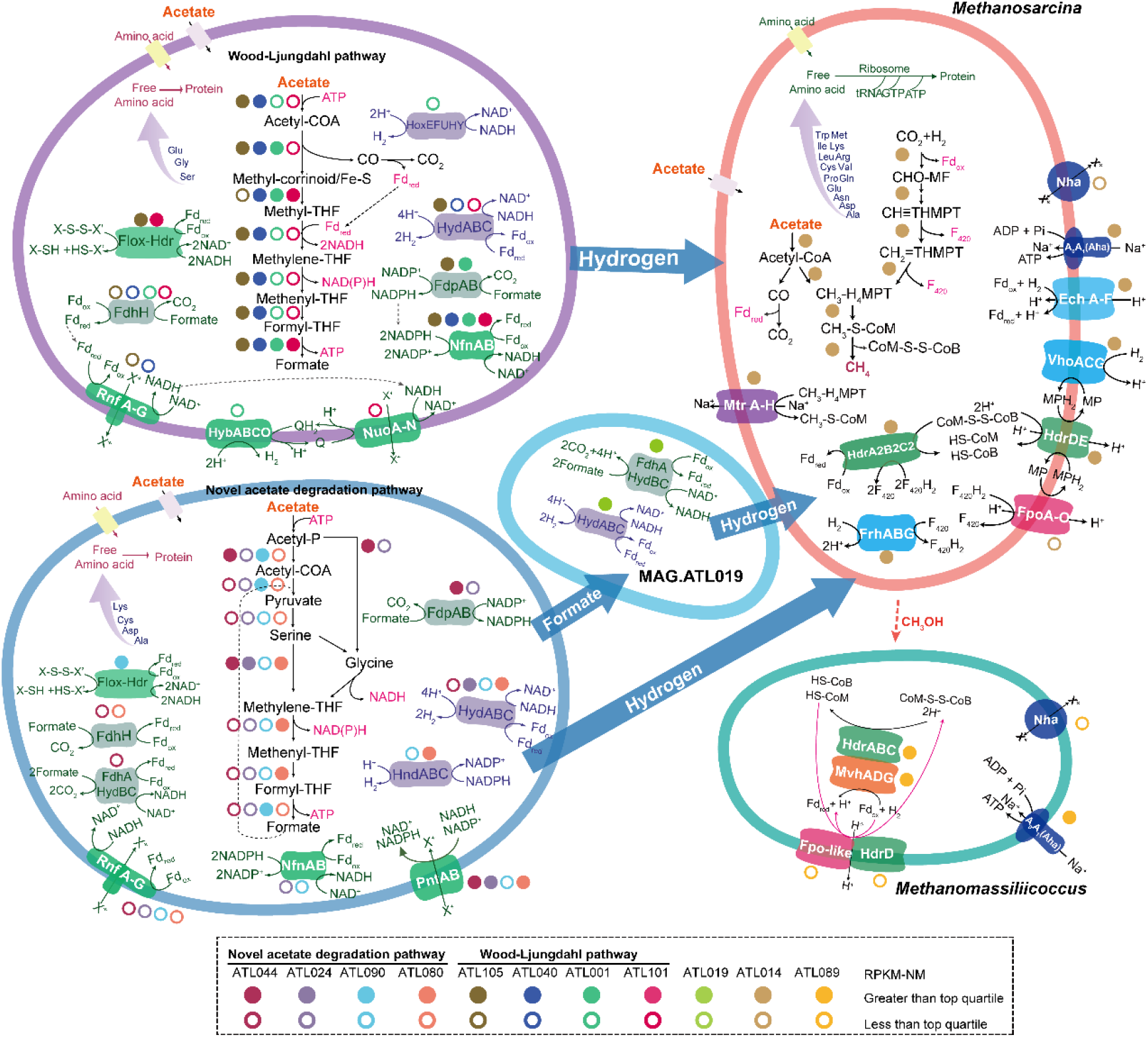
Overview of the metabolism of syntrophs and methanogens from thermophilic acetate-degrading communities. Hydrogenase, formate dehydrogenases and energy conservation pathways are abbreviated as shown in Supporting Information Tables S3-S6.

As discussed above, reducing equivalents (*i.e*., NADH, NADPH and reduced ferredoxin [Fd_red_]) involve in the actions of hydrogenases and formate dehydrogenases. Acetate oxidation via reverse WL pathway and glycine-mediated pathway generate both NADH and NADPH, and the latter pathway yields one mol of ATP per mol acetate oxidized (Fig. 2A) [17, 45]. However, Fd_red_ is not directly generated from acetate degradation. To complete acetate degradation, SAOB in acetate-degrading community encode redox complexes supporting electron transfer between (i) NAD(H) and Fd (*Rhodobacter* nitrogen fixation complex Rnf; NADH:Fd oxidoreductase; MAG.ATL080, MAG.ATL024, MAG.ATL044, MAG.ATL090, MAG.ATL105 and MAG.ATL040) [46], (ii) NADP(H) and NAD(H) and Fd (NADH-dependent NADP:Fd oxidoreductase NfnAB; MAG.ATL024, MAG.ATL090, MAG.ATL105, MAG.ATL040, MAG.ATL001 and MAG.ATL101) [17, 43, 47], (iii) NAD(H) and NADP(H) (transhydrogenase PntAB; MAG.ATL080, MAG.ATL024, MAG.ATL044 and MAG.ATL090) [48, 49], (iv) Fd and unknown electron carriers (uncharacterized oxidoreductase Flox:Hdr; MAG.ATL090, MAG.ATL105 and MAG.ATL101) [17, 45, 50] (Figs. 3A and 5; Table S6). Using these complexes, the syntrophic acetate oxidizers may employ reverse electron transport and electron bifurcation to generate H_2_.

In addition, we found that MAG.ATL044 contained a more energy-efficient pathway, which might be the reason why it possessed high abundance and activity in ATL (Figs. 3A and 5; Table S6). Analyses of metatranscriptomic indicates that MAG.ATL044 did not express H_2_ generation activity, suggesting it converted acetate to formate but not further to H_2_ and CO_2_ (Figs. 3A and 5; Table S6). This observation is consistent with previous study that some novel acetate-degrading species expressing the glycine-mediated pathway just oxidized acetate to formate (no H_2_ generation) in full-scale anaerobic digesters [51]. Formate was not detected in ATL, indicating that acetate was completely oxidized without accumulation of this metabolic intermediate. In addition, we did not detect F_420_-reducing formate oxidation (FdhAB) in *Methanosarcina* in ATL. Thus, we suspected other syntrophic species may convert the acetate-derived formate to H_2_. In agreement, metagenomics analyses showed that Clostridia (MAG.ATL106) and Anaerolineaceae (MAG.ATL019) highly expressed formate dehydrogenases and hydrogenases (Fig. 5 and Table S6), suggesting potential involvement in formate oxidation [52, 53]. In contrast, another acetate degrader MAG.ATL001, which was the second most active bacterial MAG (accounting for 13.1% of the activity) highly expressed formate dehydrogenases and hydrogenase at similar levels, oxidizing acetate to H_2_ and CO_2_ via reverse WL pathway (Figs. 3A and 5; Table S6). This result suggests that acetate oxidation by MAG.ATL001 may prefer H_2_ for interspecies electron transfer in our thermophilic community. The glycine-mediated pathway avoids endergonic 5-methyl-THF oxidation, generating a yield of 1 ATP per acetate. In comparison, the reverse WL pathway hold a puzzling theoretical yield of 0 ATP per acetate [17] (Fig. 2A). This optimal energy generation strategy might partially explain the high abundance and activity of MAG.ATL044.

In order to improve our understanding of energy conservation revolving around SAO process, we compared electron transfer mechanisms of our novel enriched acetate oxidizers with previously known SAOB. We found a new formate dehydrogenase, FdpAB, located in the novel acetate oxidizers in ATL (Figs. 3A and S8; Tables S6 and S7). *T. phaeum* and *S. schinkii* encode membrane-bound cytochrome b-linked quinone-dependent formate dehydrogenase (FdnGHI) associated with proton extrusion, which were not found in novel acetate oxidizers in ATL. In addition, all the known groups of formate dehydrogenases were not identified in *T. acetatoxydans* and *P. lettingae*, though they were capable of converting acetate into CO_2_ and H_2_. These analyses implied potentially high diversity of formate dehydrogenase in SAOB. In terms of hydrogenase, electron-confurcating hydrogenases HydABC has been found in all novel acetate oxidizers and the known SAOB (except *S. ultunense*) (Figs. 3A and S8; Tables S6 and S7), suggesting such electron confurcation mechanism is universal in SAOB. However, several novel acetate oxidizers had depressed expression of HydABC. We speculate that these populations may have unknown hydrogenase and/or they may transfer formate to other syntrophic populations. Morever, Rnf, NfnAB, PntAB and HydABC were well conserved amongst some novel acetate oxidizers and known SAOB, suggesting that they may serve a core function to SAO.

In addition to interspecies electron transfer via hydrogen and formate, DIET has been conceived as a potential mechanism in extracellular electron transfer [54], which depends on electrically conductive type IV pili and outer-surface c-type cytochromes [55]. The putative acetate metabolizers (Only Gemmatimonadetes MAG.ATL080 and Thermoanaerobacteraceae MAG.ATL024) encode a type IV pilin assembly protein (PilC) and a VirB11-like ATPase (PilB), but no genes encoding the structural protein PilA, which is associated with DIET [56], were found (Table S6). A c-type cytochrome was detected in Gemmatimonadetes (MAG.ATL080) and Anaerolineae (MAG.ATL001 and MAG.ATL101), and only MAG.ATL080 and MAG.ATL101 high expressed this gene (Table S6). Therefore, DIET may play a role in syntrophic acetate degradation, but not the essential role. The roles of DIET in these novel acetate degraders remain unclear but warrant further attention.

#### Energy**-**conserving metabolisms in methanogens

In the archaeal community, *Methanosarcina* MAG.ATL014 obtained the electrons from intermediate H_2_ through highly expression of methanophenazine(MP)-reducing hydrogenase (VhoGAC), energy-converting [NiFe] hydrogenase (EchA-F), and F_420_ reduction hydrogenase (FrhABG) (Figs. 3B and 5; Table S4). The electrons provided by FrhABG were transferred to F_420_ to produce F_420_H_2_. The electrons carried by F_420_H_2_ were used for two reduction steps (methenyl-H_4_MPT→methylene-H_4_MPT→methyl-H_4_MPT) in hydrogenotrophic methanogenesis, as well as transferred to MP via a FpoF-lacking FpoA-O. Finally, MPH_2_ reduced by VhoGAC and FpoA-O transferred electrons to CoM-S-S-CoB via heterodisulphide reductase (HdrDE) (Figs. 3B and 5; Table S4). The metabolism yields energy by forming proton motive force via HdrDE [57, 58]. *Methanosarcina* also highly expressed the heterodisulfide reductase homologous HdrA2B2C2 that oxidize F_420_H_2_ with the reduction of Fd_ox_ and CoB-S-S-CoM through flavin-based electron bifurcation [59, 60]. The H_2_/CO_2_-dependent methanogen *Methanothermobacter* (MAG.ATL045) has genes for H_2_ oxidation via reverse electron transport (EhaA-T rather than EchA-F), electron bifurcation (MvhADG-HdrABC), and F_420_ reduction (FrhABG). Nevertheless, *Methanothermobacter* did not express all of above genes for H_2_/formate oxidation (Figs. 3B and 5; Table S4). *Methanomassiliicoccus* highly expressed genes for electron-bifurcating H_2_ oxidation and a putative ferredoxin: heterodisulfide oxidoreductase complex for electron transduction from H_2_ to methanol-reducing methanogenesis (via MvhADG, HdrABC, FpoF-lacking Fpo-like, and HdrD) (Fig.3B and 5; Table S4). The high H_2_ oxidation activity detected in *Methanosarcina* MAG.ATL014 in ATL was associated with consumption of H_2_ produced from SAO and formate oxidation. Moreover, *Methanosarcina* higher expressed the genes involved in CO_2_-reducing pathway in ATL compared to PTL and VTL (Tables S3 and S4, [28, 30]). These results implied that *Methanosarcina* played a multi-trophic functional role in thermophilic chemostats.

### Syntrophic metabolism and energy conservation of acetate-degrading community in propionate**-**, butyrate**-**, and isovalerate**-**fed chemostats

As acetate is an important by-product from syntrophic fatty acid degradation (*e.g.*, propionate, butyrate, and isovalerate), SAOB should be also present in the chemostats fed with these fatty acids. To investigate whether the observed novel SAOB in ATL also play roles in oxidation of other fatty acids, we analyzed the metabolic features of the MAGs involved in the microbial communities in other seven chemostats. Similar with the result in ATL, the previously known SAOB (*i.e.*, *T. phaeum*, *P. lettingae*, *S. ultunense*, *S. schinkii*, and *T. acetatoxydans*) were not detected in PTL, whereas populations related to the known acetate-oxidizing genus *Tepidanaerobacter* (MAG.BTL055 and MAG.VTL084) only displayed low activity and did not express acetate oxidizing activity in butyrate- and isovalerate-degrading communities (Fig. S7; Tables S5 and S6). In propionate-, butyrate- and isovalerate-degrading communities, we identified multiple bacterial populations associated with uncultured Clostridia, Thermoanaerobacteraceae, Anaerolineae, Gemmatimonadetes and *Thermodesulfovibrio* that encode the reverse WL and glycine-mediated pathways, associated with complementary NADPH re-oxidation and H_2_/formate-generating enzymes (Fig.S7; Tables S5 and S6). Based on the above criteria (See Results and discussion 3.4.1 for details of the settings), among these populations, *Thermodesulfovibrio* (MAG.PTL017 and MAG.VTL073), *Desulfotomaculum* (MAG.BTL007), Thermoanaerobacteraceae populations (MAG.VTL038 and MAG.BTL014), *Clostridia* populations (MAG.PTL141 MAG.BTL065 and MAG.VTL024), and Gemmatimonadetes (MAG.BTL079 and MAG.VTL039) may syntrophically degrade acetate (the former three expressed the reverse Wood-Ljungdahl pathway and the last expressed the *Thermotogae*-associated pathway; Fig.6; Tables S5 and S6). These results confirmed our previous study, in which members of Clostridia, Thermoanaerobacteraceae, Anaerolineae, and *Thermodesulfovibrio* were labeled with ^13^C_2_-sodium acetate in DNA stable isotope probing assays [23]. Therefore, these microorganisms were potential acetate degrader.

**Fig. 6.**
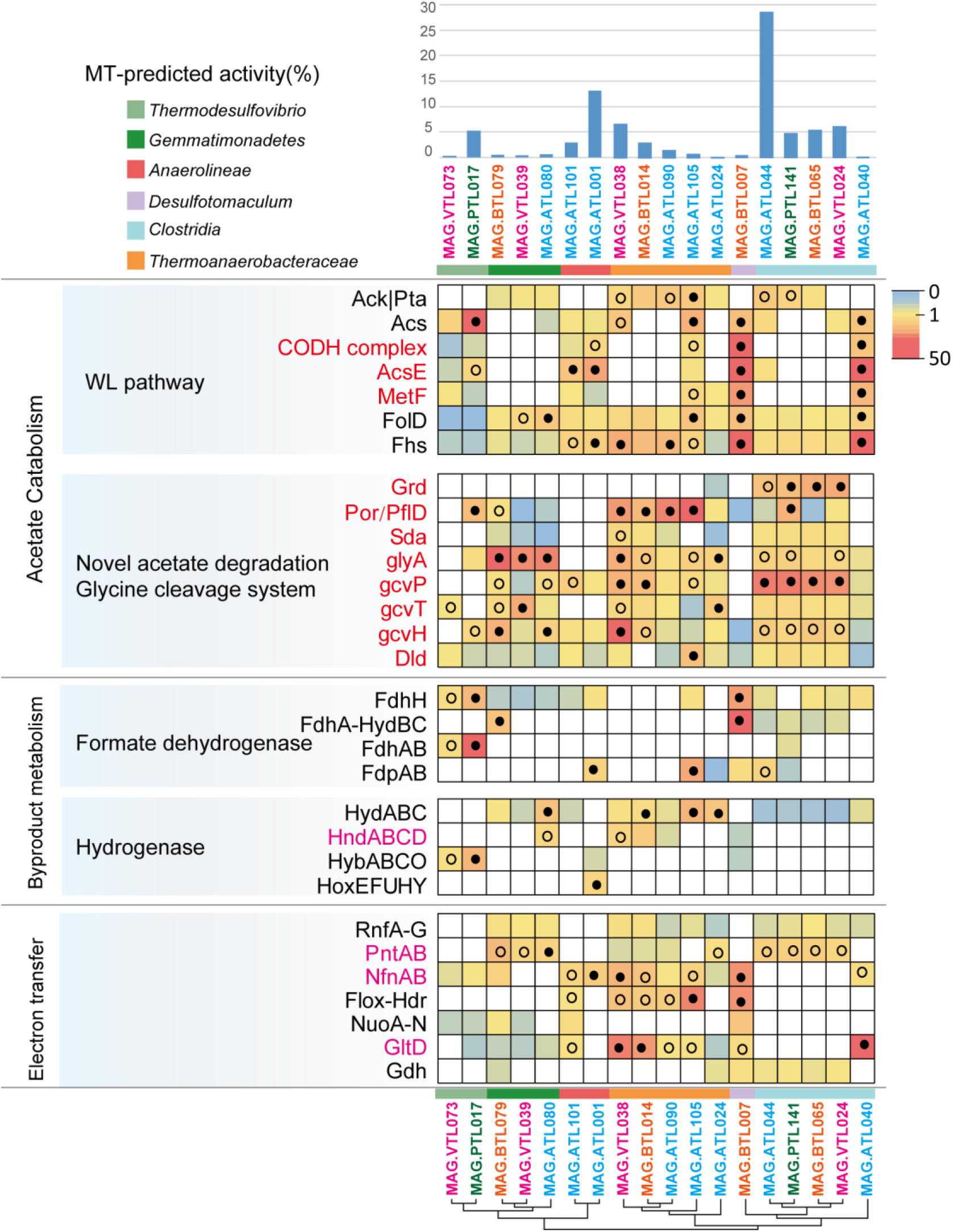
Gene expression level for acetate oxidation, H_2_/formate metabolism, and electron transfer genes in syntrophs which may syntrophically degrade acetate from thermophilic chemostats. For each MAG, the percentages of the metatranscriptomic (MT) reads mapped to the MAG out of the metatranscriptomics mapped to all MAGs (both *Bacteria* and *Archaea*) are shown. The gene expression levels are calculated as reads per kilobase of transcript per million reads mapped to individual MAG (RPKM) normalized to the median gene expression for the corresponding MAG (RPKM-NM) averaged from duplicate samples. Pathways containing genes with RPKM-NM greater than the octile and quartile are marked (filled and open dots, respectively). Enzyme abbreviations and their corresponding genes are elaborated in Supporting Information Tables S5-S6.

### Biosynthetic metabolism of acetate oxidizers

In our chemostats, the novel SAOB used acetate as the carbon and energy source for producing H_2_, reducing equivalents (*i.e*., NADH, NADPH and reduced ferredoxin [Fd_red_]), and ATP, which provided the cell with energy for biomass biosynthesis. We further found that these novel acetate degraders encode pathways for converting acetyl-CoA to pyruvate, as well as other important precursors for biosynthesis of sugars, amino acids (AAs) and nucleotides, and pathways for AAs degradation (Tables S8-S10).

In complex microbial communities, microbes are frequently observed to interact other individuals by exchanging the above mentioned metabolites as public goods [61, 62], we thus set out to test whether this is also the case in our novel SAOB. Strikingly, we found that no single SAOB contains all the genes encoding the synthesis of an entire suite of amino acids (AAs) (Fig. S9 and Table S8), suggesting that the exchange of the essential AAs (one typical set of public goods reported previously [63, 64]) is common between these novel SAOB in thermophilic fatty acid-degrading community. Further analyses indicated that acetate degraders tended to lose/ lowly express the AA biosynthetic pathway with higher energy cost (Fig. S9 and Table S8). In addition, the populations which closely related to each other possessed similar capabilities for AA biosynthesis. The Clostridia-related acetate metabolizers (MAG.ATL044, MAG.PTL141, MAG.BTL065 and MAG.VTL024), which displayed the dominant activity in individual chemostat and proposed glycine-mediated acetate oxidation pathway, only synthesized three AAs (*i.e.*, glutamate, glycine, and serine) (Fig. S9 and Table S8). In comparison with the most active acetate degraders (MAG.ATL044, MAG.PTL141, MAG.BTL065 and MAG.VTL024), other acetate degraders have genes for synthesizing more AAs, whereas they had depressed expression of biosynthesis for the majority of AAs (Fig. S9 and Table S8). According to the Black Queen Hypothesis, when an individual loses an energy-expensive function, it becomes a ‘beneficiary’, who scavenges the public goods, such as AAs, from other individuals (helpers) for survival [65, 66]. The ‘beneficiary’ strain will expand in the community until the production of public goods, such as AAs, is just sufficient to support the balanced community. AA biosynthesis is an energy-consuming process, and therefore, the acetate metabolizers can invest more energy to their metabolism on acetate oxidation rather than biosynthesis. These results suggest that the metabolic exchange of public goods between SAOB and other members plays a significant role on maintaining high efficiency of acetate metabolism during the AD process.

### The prevalence of the novel SAOB across diverse thermophilic AD communities

The novel acetate degraders from uncultured lineages in this study (members of Clostridia, Thermoanaerobacteraceae, Anaerolineae, and Gemmatimonadetes) from ATL, PTL, BTL and VTL were distantly related to any isolated species (Fig. S10). Moreover, these novel acetate oxidizers were closely related (≥97% similarity; Fig. S11) to uncultured populations detected in thermophilic anaerobic digesters feeding with a wide diversity of substrates (*e.g.*, sole carbon source [67, 68], binary carbon source [69], municipal waste [70, 71], agricultural waste [72, 73] and industrial waste [74, 75]). Therefore, the novel syntrophic lineages revealed in the present study are the universal and probably core species that play a critical role in thermophilic methanogenic process in thermophilic anaerobic digestion.

## Conclusion

In summary, we combined metagenomics and metatrascriptomics to characterize novel syntrophic acetate oxidizers, including Clostridia, Thermoanaerobacteraceae, Anaerolineae, Gemmatimonadetes and *Thermodesulfovibrio* members, in thermophilic anaerobic chemostats. The high expression of genes involved in acetate oxidation and energy conservation systems indicated that these acetate-oxidizing species played an essential role in syntrophic acetate degradation. *Methanosarcina thermophile* highly performed acetate and H_2_ utilizing methanogenesis, consuming H_2_ derived from acetate oxidizers. These findings improve our understanding of the phylogenetic diversity and metabolic characteristics of the syntrophic acetate oxidizers and their interaction with methanogens, although thoroughly characterization of these newly proposed acetate oxidizers still requires further effortful cultivation-based studies. Managing the performance of these novel bacterial clades may be key to improve the efficiency of anaerobic digestion, facing against serious challenges of the environment degradation and energy shortage.

## Materials and Methods

### Operation of thermophilic anaerobic chemostats

Eight thermophilic (55 °C) anaerobic chemostats (Fig. S12) were constructed using continuous stirred tank reactors (CSTRs), each with a working volume of 1.8 L. The seed sludge for inoculating acetate- and propionate-degrading chemostats was obtained from a thermophilic anaerobic digester treating kitchen waste (Sichuan Province, China), and the seed sludge for inoculating butyrate- and isovalerate-degrading chemostats was from a swine manure treatment plant (Sichuan Province, China) (Table S1). The seed sludge was rinsed with the washing solution (synthetic wastewater without carbon sources) thrice under anaerobic condition, diluted to 1.8 L, and then inoculated into each chemostat.

The thermophilic chemostats were fed with synthetic wastewater containing acetate, propionate, butyrate, or isovalerate as the sole carbon source, respectively (total organic carbon = 8000 mg·L^-1^) (Table S1). The synthetic wastewater contains 0.3 g/L KH_2_PO_4_, 4.0 g/L KHCO_3_, 1.0 g/L NH_4_Cl, 0.6 g/L NaCl, 0.82 g/L MgCl_2_·6H_2_O, 0.08 g/L CaCl_2_·2H_2_O, 0.1 g/L cysteine-HCl·H_2_O; 10 mL trace element solution containing 21.3 mg·L^-1^ NiCl_2_·6H_2_O and 24.7 mg·L^-1^ CoCl_2_·6H_2_O; and 10 mL vitamin solution. 5.46 g sodium acetate and 16.0 g acetic acid were added for acetate synthetic wastewater, 4.27 g sodium propionate and 13.16 g propionic acid were added for propionate synthetic wastewater, 14.67 g butyric acid and 1.33 g NaOH were added for butyrate synthetic wastewater, 13.60 g isovalerate acid and 1.07 g NaOH were added for isovalerate synthetic wastewater.

Briefly, the thermophilic chemostats were incubated in thermostat-controlled water-baths (TR-2A, ASONE, Osaka, Japan). Broth in each chemostat was thoroughly mixed using a magnetic stirrer at 200–300 rpm. Synthetic wastewater was supplied to the chemostats by continuous feeding under an atmosphere of N_2_, and the effluent over flowed automatically via a U-type tube. Biogas was collected with a gas holder. Initially, two replicated chemostats were performed for each carbon source, so a total of eight chemostats were operated. The eight thermophilic chemostats were initially operated at a dilution rate of 0.01 d^−1^ and then increased to 0.025 d^−1^ (hydraulic retention time [HRT] = 40 d), after a period of operation at a dilution rate of 0.025 d^−1^, dilution rates of four chemostats were increased to 0.05 d^−1^ (hydraulic retention time [HRT] = 20 d). For simplicity, we used A (acetate), P (propionate), B (butyrate), and V (isovalerate) to denote four different carbon sources used in these chemostats; L was used to indicate the lower dilution rate (0.025 d^−1^), whereas H indicated the higher dilution rate (0.05 d^− 1^). Therefore, the eight chemostats were named as ATL, PTL, BTL, VTL, ATH, PTH, BTH, and VTH, respectively (Table S1).

Broth was sampled every week for fluorescence microscopic observation and physicochemical analyses of pH, suspended solids (SS), volatile suspended solids (VSS), TOC, as well as volatile fatty acids (VFAs) using the same protocols described previously [28]. In addition, the methane and H_2_ contents in biogas were determined by a gas chromatograph (GC-2014C, Shimadzu, Kyoto, Japan). During the steady operation period, biomass collected from broth was used for DNA and RNA extraction.

### DNA and RNA extraction, 16S rRNA gene PCR, sequencing, and data processing

Sludge was collected from 40 mL broth from each chemostat (ATL, day 286; ATH, day 258; PTL, day 283; PTH, day 400; BTL, day 284; BTH, day 398; VTL, day 300; VTH day 360) by centrifugation at 13000 ×g at 4℃ for 10 min. The sludge was rinsed thrice with sterile phosphate buffer saline (PBS) (10 mM, pH 7.5). Total DNA and RNA (duplicates for each sample) were extracted via cyltrimethyl ammonium bromide (CTAB) method [76]. Total RNA was reverse transcribed using PrimeScript^TM^ RT reagent Kit with gDNA Eraser (Perfect Real Time) according to the manufacturer’s protocol (Takara, Kusatsu, Japan). DNA and cDNA samples were subjected to 16S rRNA gene amplicon sequencing. The 16S rRNA genes of both bacteria and archaea were amplified through PCR using universal primers 515F (5′-GTGCCAGCMGCCGCGGTAA-3′) and 909R (5′-CCCCGYCAATTCMTTTRAGT-3′) targeting the V4-V5 hypervariable regions. The 16S rRNA gene amplicon sequencing was performed on an Illumina MiSeq platform (Illumina, San Diego, CA, USA) according to the standard protocols by Majorbio Bio-Pharm Technology Co. Ltd. (Shanghai, China). The data processing was conducted using the protocol as previously reported [23].

### Metagenomic and metatranscriptomic sequencing, as well as the associated bioinformatics analyses

Sludge samples for metagenomic sequencing were collected on different dates from ATL (day 306 and 307), PTL (day 223, 293 and 318), BTL (day 251 and 252), VTL (day 295 and 296). Similarly, sludge samples for metatranscriptomic sequencing were collected on two different dates from ATL (day 302 and 305), PTL (day 223, and 293), BTL (day 247 and 249), VTL (day 291 and 293). Total DNA and RNA were extracted using CTAB method [76]. Metagenomic DNA was sequenced on an Illumina HiSeq 2000 platform (Illumina). The paired-end reads (2 × 150 bp) were trimmed via Trimmomatic v0.36 [77] with a quality cutoff of 30, sliding window of 6 bp and minimum length cutoff of 100 bp. The clean reads from different metagenomes of the same chemostat were co-assembled via SPAdes v.3.5.0 [78], binned through MetaBAT [79], and checked for completeness and contamination using CheckM [80]. The completeness was calculated based on the number of expected marker genes present in MAGs, while the contamination based on the number of expected marker genes present in multiple copies. Genes were annotated using Prokka [81] and manual curation was performed as described previously [30]. Phylogenomic trees were built with PhyloPhlAn v0.99 (“-u” option) [82], and the tree was edited using iTOL [83].

For metatranscriptomics sequencing, total RNA was purified by removing residual DNA by an RNase-free DNase set (Qiagen, Hilden, Germany). Ribosomal RNA was removed from the DNase-treated RNA via the Ribo-Zero rRNA Removal Kits (Illumina). RNAseq libraries were constructed using the TruSeq RNA sample prep kit (Illumina) with the standard protocol. The libraries were sequenced on an Illumina HiSeq2000 sequencer. The paired-end (2 × 150 bp) metatranscriptomic reads were trimmed as DNA-trimming step described above and mapped to MAGs using the BBMap with the parameters as: minid = 1 (v35.85; http://sourceforge.net/projects/bbmap/). The gene expression levels of functional genes from each MAG were calculated as reads per kilobase transcript per million reads mapped to the MAG (RPKM) averaged from duplicate samples. The RPKM was further normalized to the median gene expression level in the heat map illustration for each bin (RPKM-NM) averaged from duplicate samples [33]. Raw sequence data reported in this paper have been deposited (PRJCA005330) in the Genome Sequence Archive in the BIG Data Center, Chinese Academy of Sciences under accession codes CRA004311 for 16S rRNA gene, metagenomics and metatranscriptomics sequencing data that are publicly accessible at http://bigd.big.ac.cn/gsa.

## Acknowledgements

This study was funded by the National Natural Science Foundation of China (No. 51678378) and the Ministry of Science and Technology of China (No. 2016YFE0127700).

## Notes

### Competing Interest Statement

The authors have declared no competing interest.

