## Supporting Information for "Chasing the metabolism of novel syntrophic acetate-oxidizing bacteria in thermophilic methanogenic chemostats"

***for***

**Supplementary figures**

**-Fig. S1.** Performance of the eight thermophilic chemostats. (A) ATL, 0.025 d^-1^; (B) ATH, 0.05 d^-1^; (C) PTL, 0.025 d^-1^; (D) PTH, 0.05 d^-1^; (E) BTL, 0.025 d^-1^; (F) BTH, 0.05 d^-1^; (G) VTL, 0.025 d^-1^; (H) VTH, 0.05 d^-1^.

-**Fig. S2.** Relative abundance of bacterial genera based on 16S rRNA gene amplicon sequencingin the thermophilic chemostats (RNA level).

-**Fig. S3.** Distribution of bacteria and archaea in the thermophilic chemostats based on (A) 16S rRNA gene amplicon with universal primers at DNA level; (B) 16S rRNA gene amplicon with universal primers at RNA level; (C) metagenomic abundance; (D) metatransriptomic activity.

-**Fig. S4.** Relative abundance of archaea in the thermophilic methanogenic microbial communities (DNA, DNA level; RNA, RNA level).

**-Fig. S5.** Phylogenetic analyses of metagenome-assembled genomes (MAGs) of methanogens in thermophilic chemostats. The corresponding abundance of MAGs are estimated from their metagenomic coverage calculated as the percentage of metagenomics (MG) reads mapped to each MAG relative to the total reads mapped to all bacterial and archaeal MAGs. The estimated activity of MAGs in acetate-degrading chemostat are shown as the percentage of metatranscriptomic (MT) reads mapped to each MAG relative to total reads mapped to all bacterial and archaeal MAGs (T, totoal MT reads; T1, MT reads of sampling time point 1; T2, MT reads of sampling time point 2).

-**Fig. S6.** Fluorescence microscopy of microbial communities during the steady operation period in the thermophilic chemostats. (BF) bright field, (FF) fluorescence field, (BF&FF) bright field overlaid with fluorescence field.

**-Fig. S7.** Gene expression level for acetate oxidation, H_2_/formate metabolism, and electron transfer genes in syntrophs from thermophilic chemostats. For each MAG, the percentages of the metatranscriptomic (MT) reads mapped to the MAG out of the metatranscriptomics mapped to all MAGs (both *Bacteria* and *Archaea*) are shown. The gene expression levels are calculated as reads per kilobase of transcript per million reads mapped to individual MAG (RPKM) normalized to the median gene expression for the corresponding MAG (RPKM-NM) averaged from duplicate samples. Pathways containing genes with RPKM-NM greater than the octile and quartile are marked (filled and open dots, respectively). Enzyme abbreviations and their corresponding genes are elaborated in Supporting Information Tables S5-S6.

**-Fig. S8.** Energy conservation strategies of known SAOB. Enzyme abbreviations and their corresponding genes are elaborated in Supporting Information Tables S7.

**-Fig. S9.** Gene expression levels for amino acid biosynthesis (A) and degradation (B) of syntrophs in thermophilic chemostats. For each MAG, the percentages of the metatranscriptomic (MT) reads mapped to the MAG out of the metatranscriptomics mapped to all MAGs (both *Bacteria* and *Archaea*) are shown. The gene expression levels are calculated as reads per kilobase of transcript per million reads mapped to individual MAG (RPKM) normalized to the median gene expression for the corresponding MAG (RPKM-NM) averaged from duplicate samples. Pathways containing genes with RPKM-NM greater than the octile and quartile are marked (filled and open dots, respectively). Enzyme abbreviations and their corresponding genes are elaborated in Supporting Information Table S8-S9.

**-Fig. S10.** Phylogenetic analyses of MAGs of syntrophs in thermophilic chemostats.

**-Fig. S11.** Ecological characterization of the novel acetate oxidizer. The orange tiles represent environments from which sequences related closely (≥ 97% similarity) to the novel acetate oxidizer in this study.

-**Fig. S12.** Schematic diagram of the thermophilic completely stirred tank reactor.

**Supplementary tables**

**-Table S1.** The carbon source, dilute rate, and seed sludge of the thermophilic chemostats.

**-Table S2.** General characteristics of metagenome-assembled genomes (MAGs) acquired in this study.

**-Table S3.** Locus tags for genes and gene expression level (PRKM and PRKM-NM) involved in methanogenesis found in methanogens from thermophilic chemostats.

**-Table S4.** Locus tags for genes and gene expression level (PRKM and PRKM-NM) involved in energy conservation found in methanogens from thermophilic chemostats.

**-Table S5.** Locus tags for genes and gene expression level (PRKM and PRKM-NM) involved in acetate oxidation found in key syntrophs from thermophilic chemostats.

**-Table S6.** Locus tags for genes and gene expression level (PRKM and PRKM-NM) involved in H_2_/formate generation and electron transfer found in key syntrophs from thermophilic chemostats.

**-Table S7.** Locus tags for genes involved in H_2_/formate generation and electron transfer found in known SAOB.

**-Table S8.** Locus tags for genes and gene expression level (PRKM and PRKM-NM) involved in amino acid biosynthesis found in MAGs from thermophilic chemostats in this study, and locus tags for genes involved in amino acid biosynthesis found in two model homoacetogens and five known syntrophic acetate oxidizers.

**-Table S9.** Locus tags for genes and gene expression level (PRKM and PRKM-NM) involved in amino acid degradation found in MAGs from thermophilic chemostats in this study, and locus tags for genes involved in amino acid degradation found in two model homoacetogens and five known syntrophic acetate oxidizers.

**-Table S10.** Locus tags for genes and gene expression level (PRKM and PRKM-NM) involved in key metabolisms found in MAGs from thermophilic chemostats in this study.

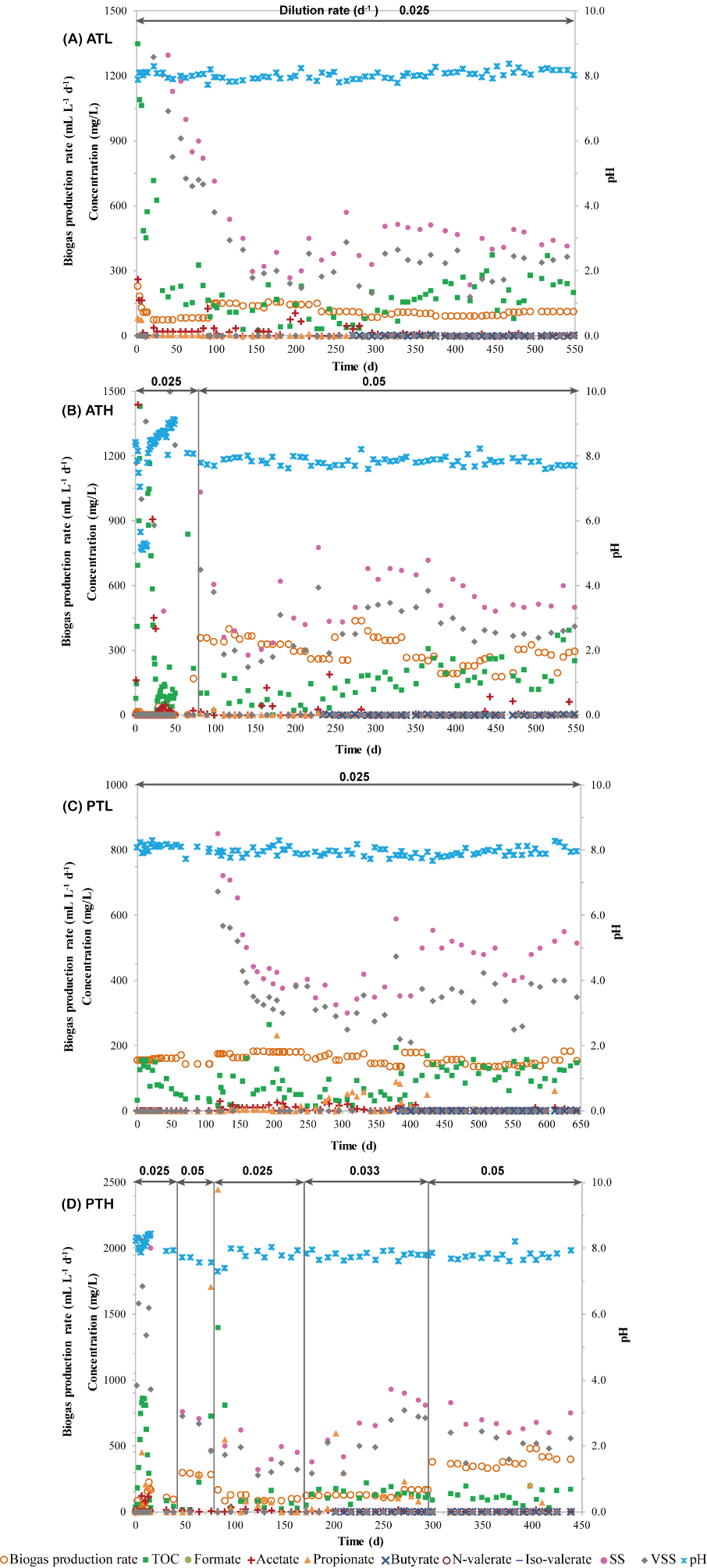

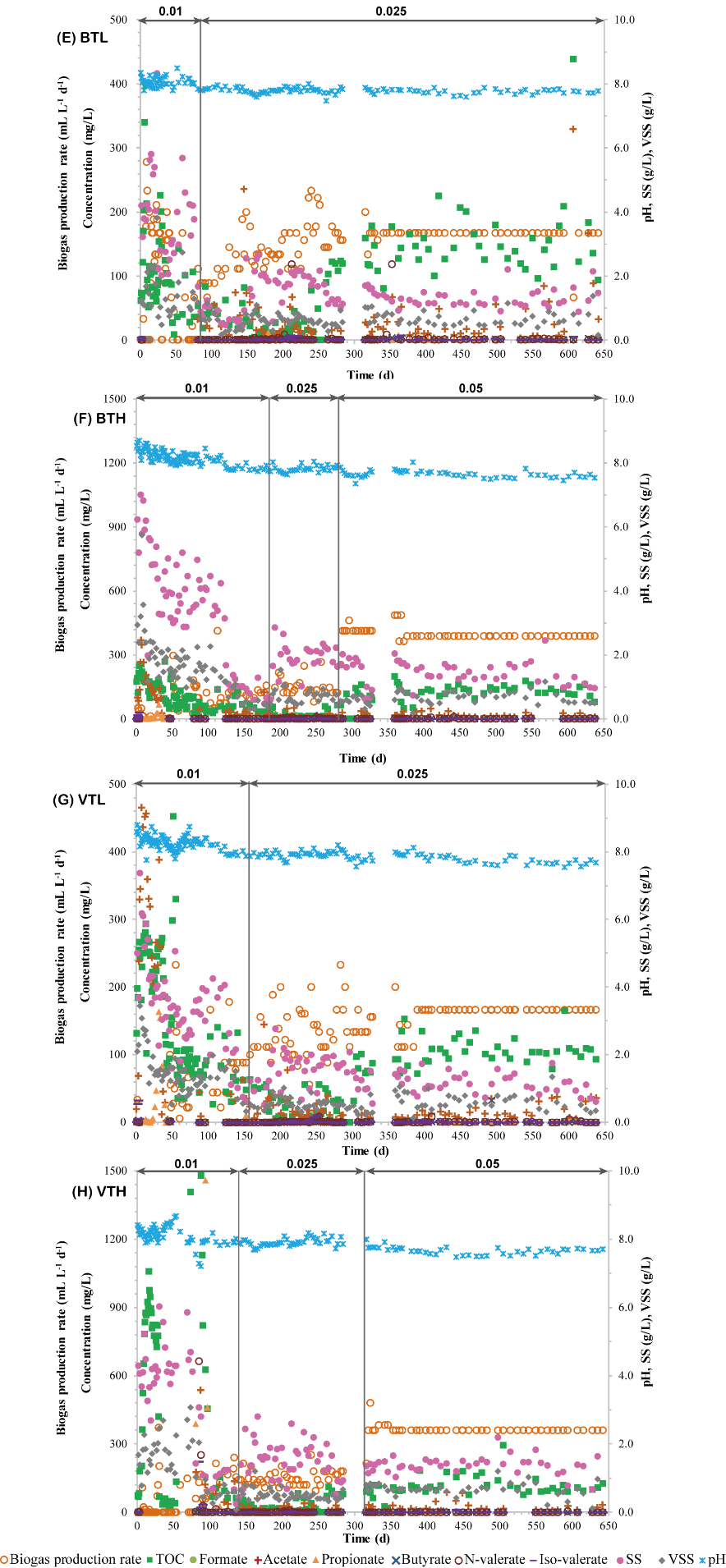

**Fig. S1.** Performance of the eight thermophilic chemostats. (A) ATL, 0.025 d^-1^; (B) ATH, 0.05 d^-1^; (C) PTL, 0.025 d^-1^; (D) PTH, 0.05 d^-1^; (E) BTL, 0.025 d^-1^; (F) BTH, 0.05 d^-1^; (G) VTL, 0.025 d^-1^; (H) VTH, 0.05 d^-1^.

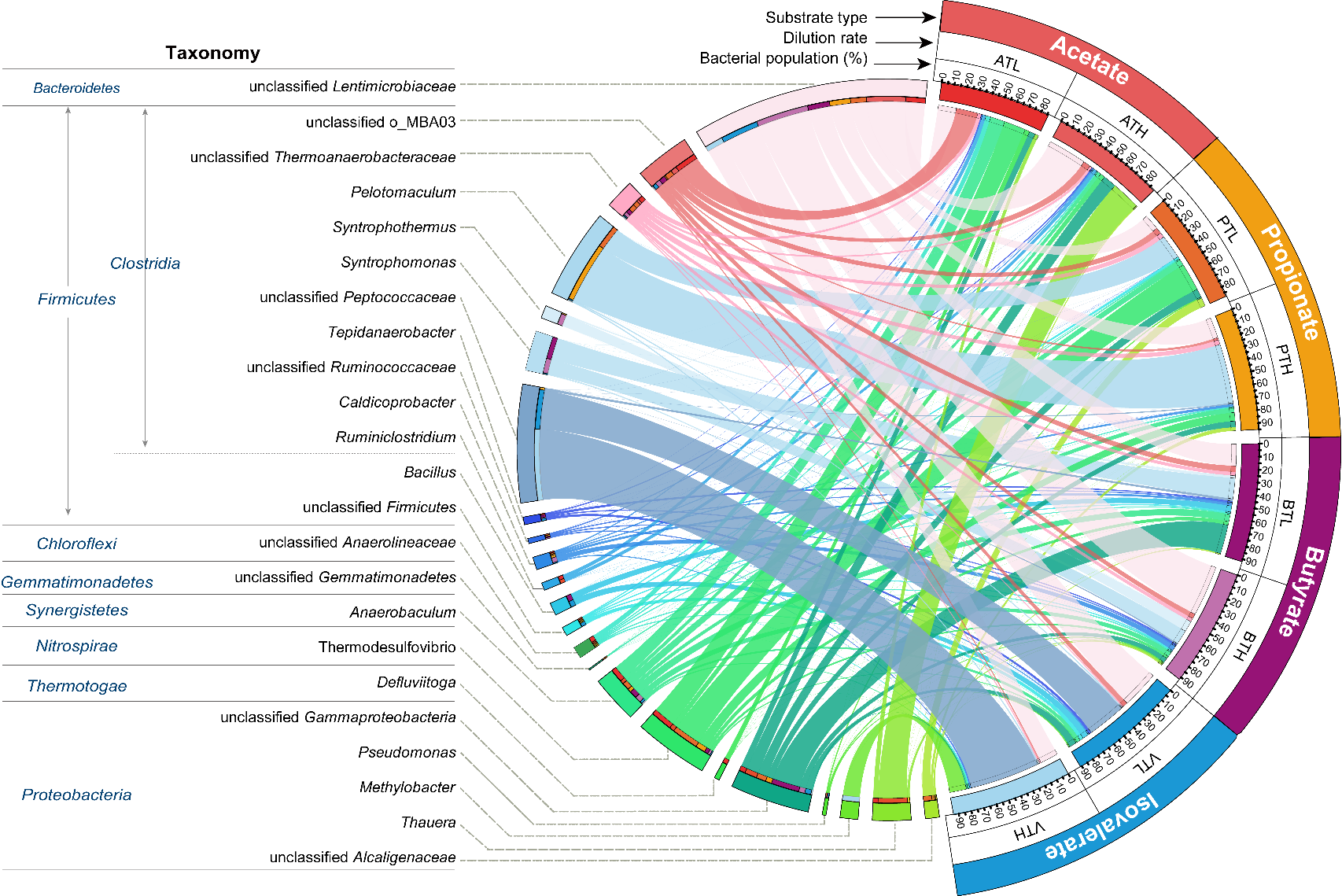

**Fig. S2.** Relative abundance of bacterial genera based on 16S rRNA gene amplicon sequencingin the thermophilic chemostats (RNA level).

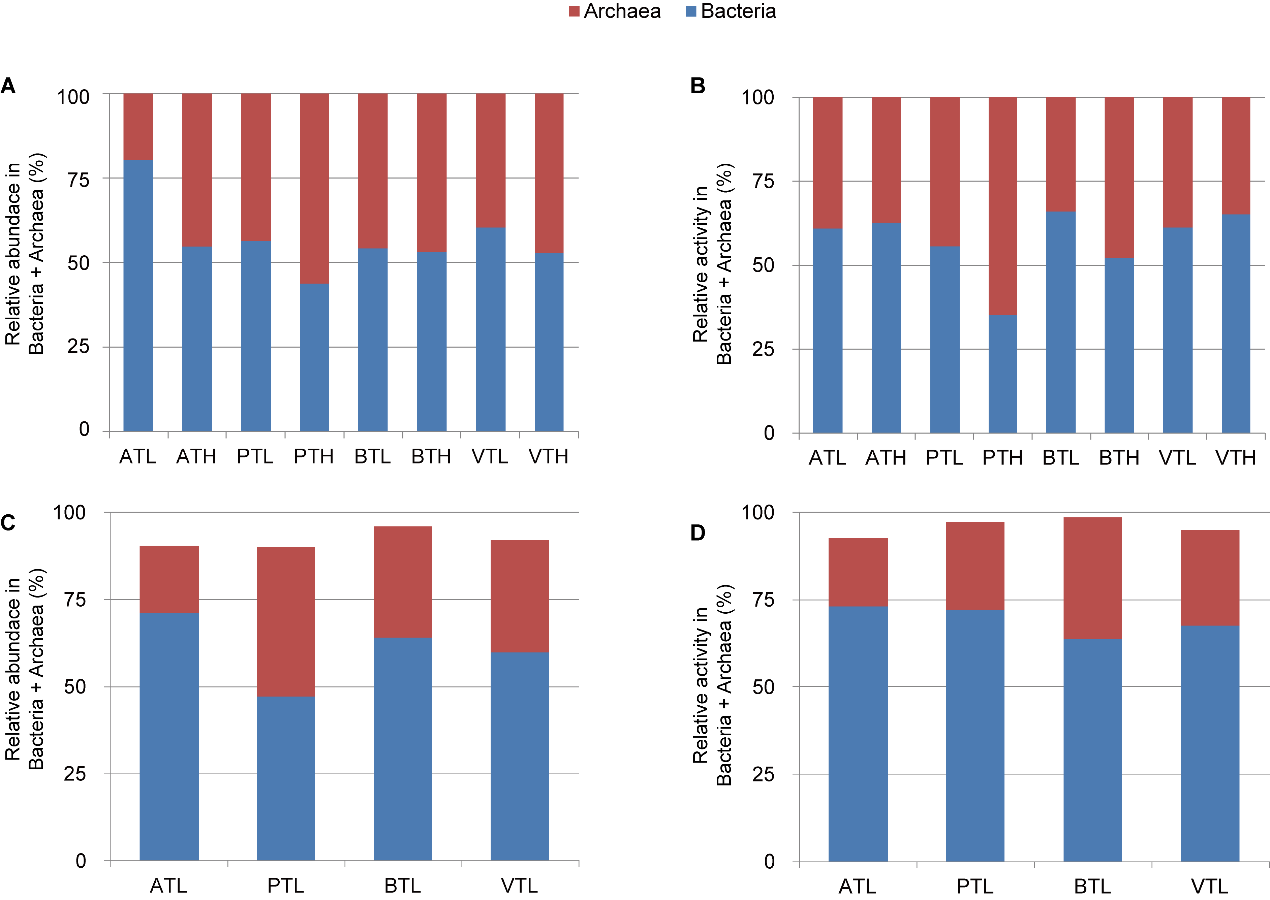

**Fig. S3.** Distribution of Bacteria and Archaea in the thermophilic chemostats based on (A) 16S rRNA gene amplicon with universal primers at DNA level; (B) 16S rRNA gene amplicon with universal primers at RNA level; (C) metagenomic abundance; (D) metatransriptomic activity.

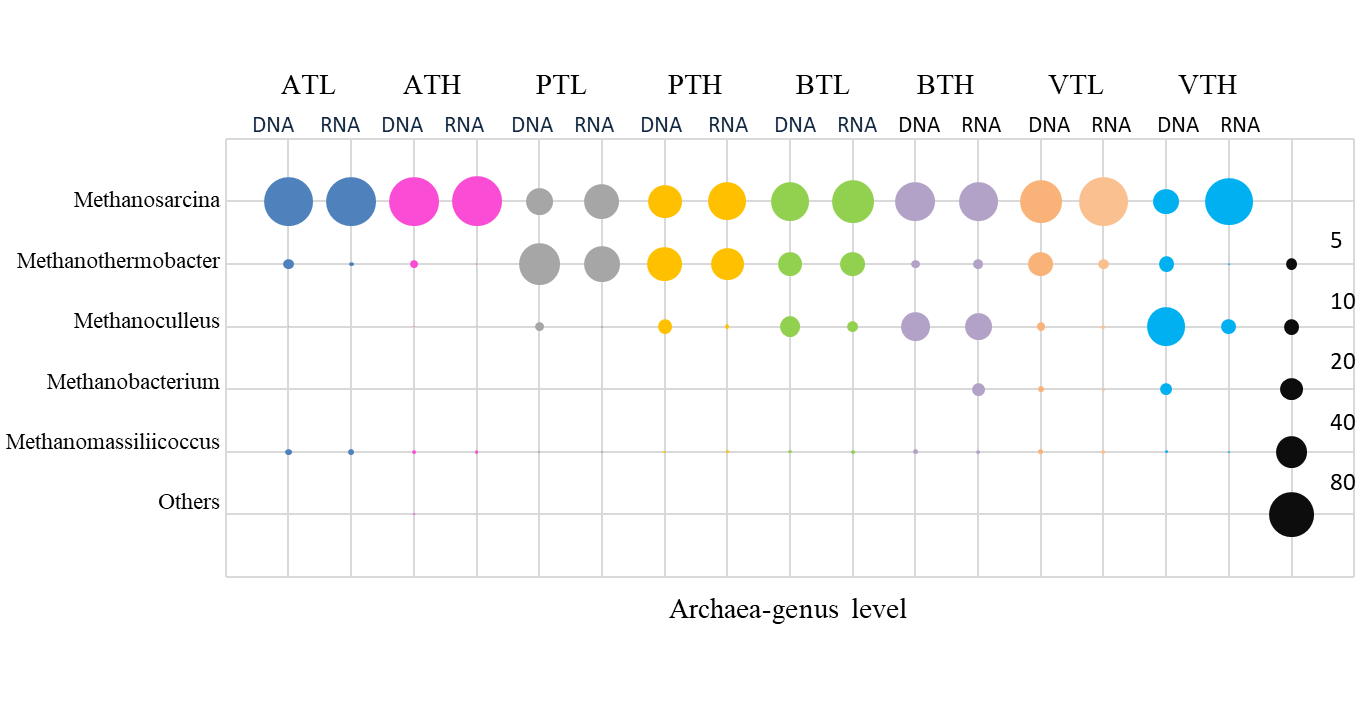

**Fig. S4.** Relative abundance of archaea in the thermophilic methanogenic microbial communities (DNA, DNA level; RNA, RNA level).

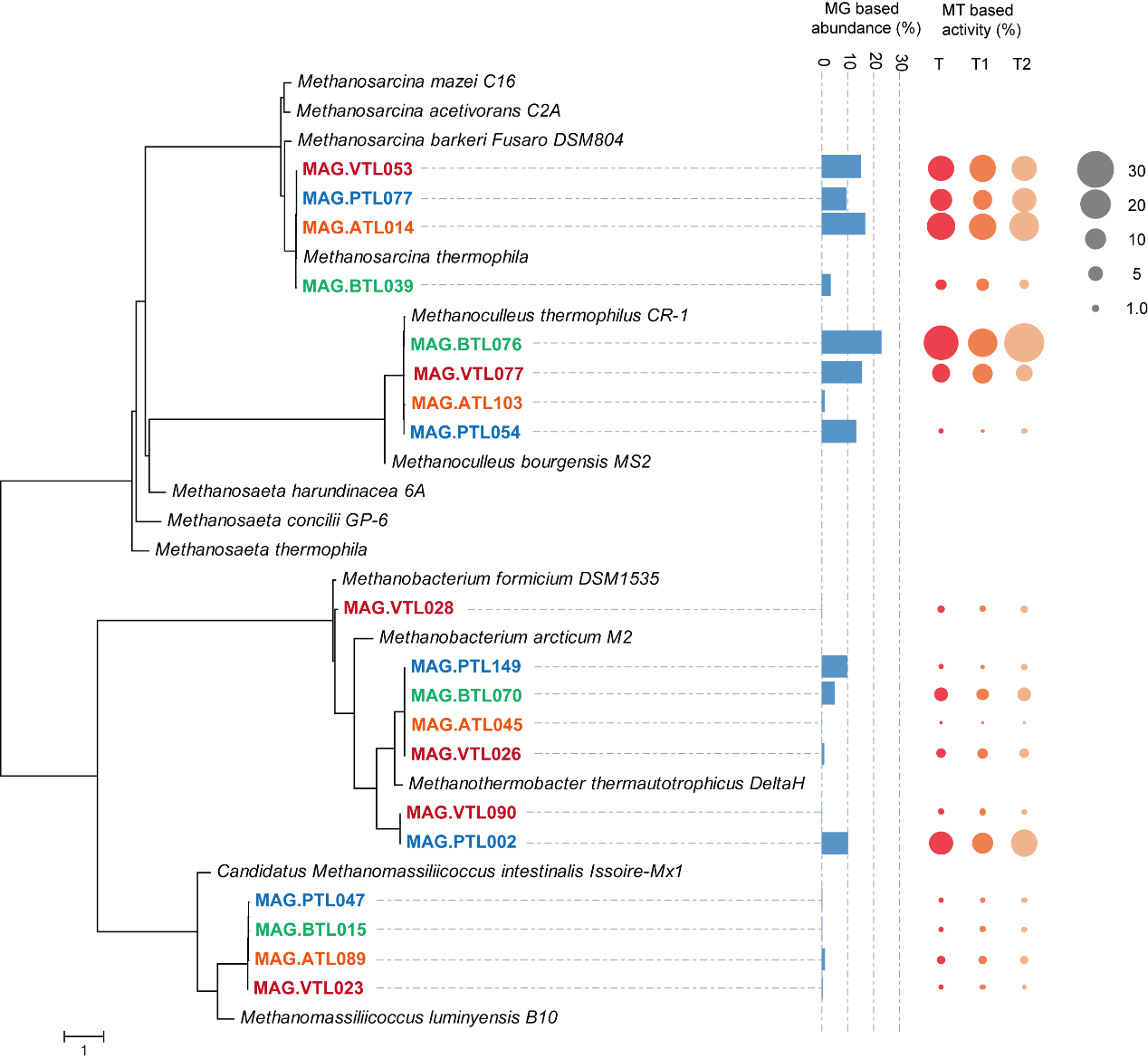

**Fig. S5.** Phylogenetic analyses of metagenome-assembled genomes (MAGs) of methanogens in thermophilic chemostats. The corresponding abundance of MAGs are estimated from their metagenomic coverage calculated as the percentage of metagenomics (MG) reads mapped to each MAG relative to the total reads mapped to all bacterial and archaeal MAGs. The estimated activity of MAGs in acetate-degrading chemostat are shown as the percentage of metatranscriptomic (MT) reads mapped to each MAG relative to total reads mapped to all bacterial and archaeal MAGs (T, totoal MT reads; T1, MT reads of sampling time point 1; T2, MT reads of sampling time point 2).

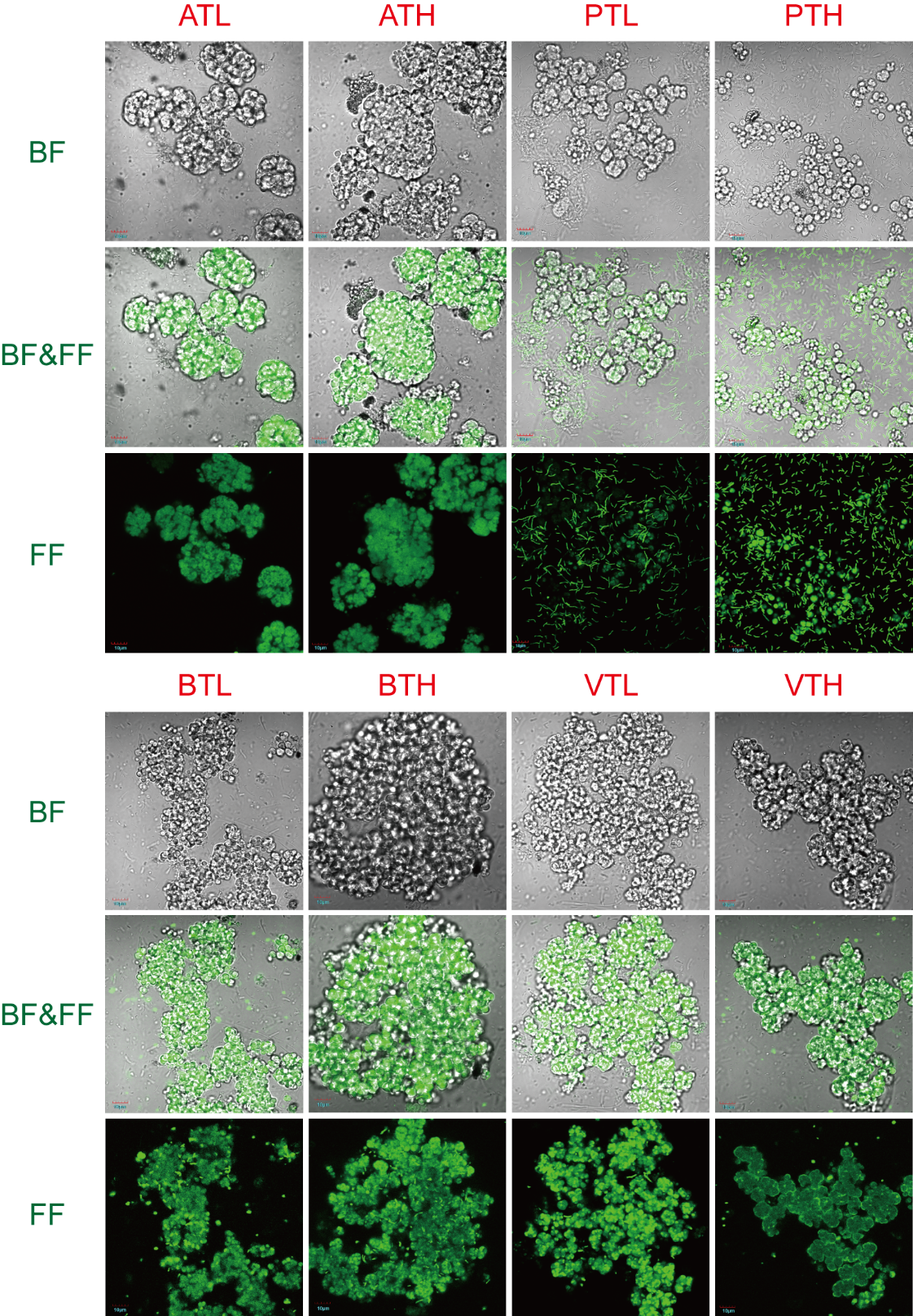

**Fig. S6.** Fluorescence microscopy of microbial communities during the steady operation period in the thermophilic chemostats. (BF) bright field, (FF) fluorescence field, (BF&FF) bright field overlaid with fluorescence field.

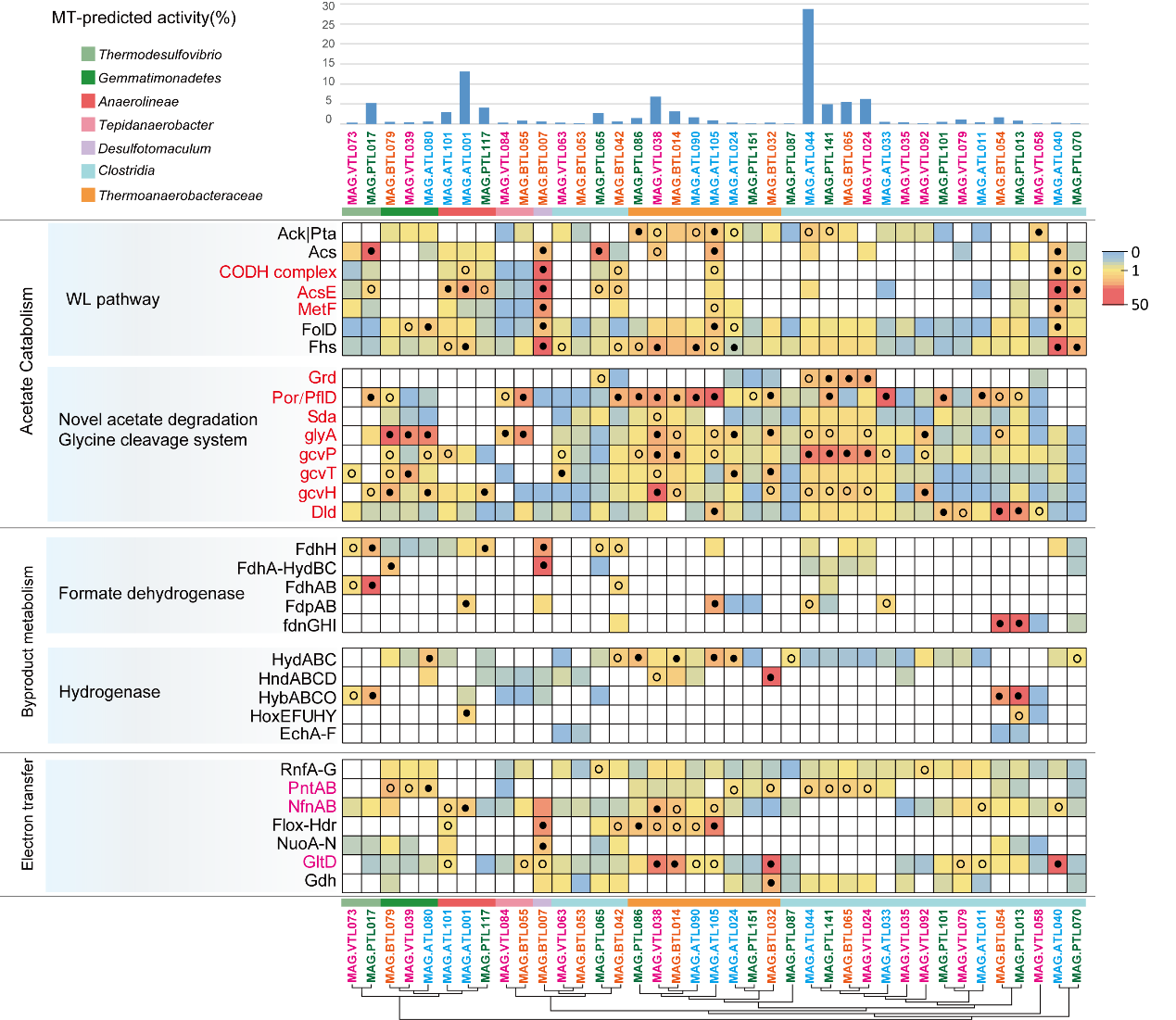

**Fig. S7.** Gene expression level for acetate oxidation, H_2_/formate metabolism, and electron transfer genes in syntrophs from thermophilic chemostats. For each MAG, the percentages of the metatranscriptomic (MT) reads mapped to the MAG out of the metatranscriptomics mapped to all MAGs (both *Bacteria* and *Archaea*) are shown. The gene expression levels are calculated as reads per kilobase of transcript per million reads mapped to individual MAG (RPKM) normalized to the median gene expression for the corresponding MAG (RPKM-NM) averaged from duplicate samples. Pathways containing genes with RPKM-NM greater than the octile and quartile are marked (filled and open dots, respectively). Enzyme abbreviations and their corresponding genes are elaborated in Supporting Information Tables S5-S6.

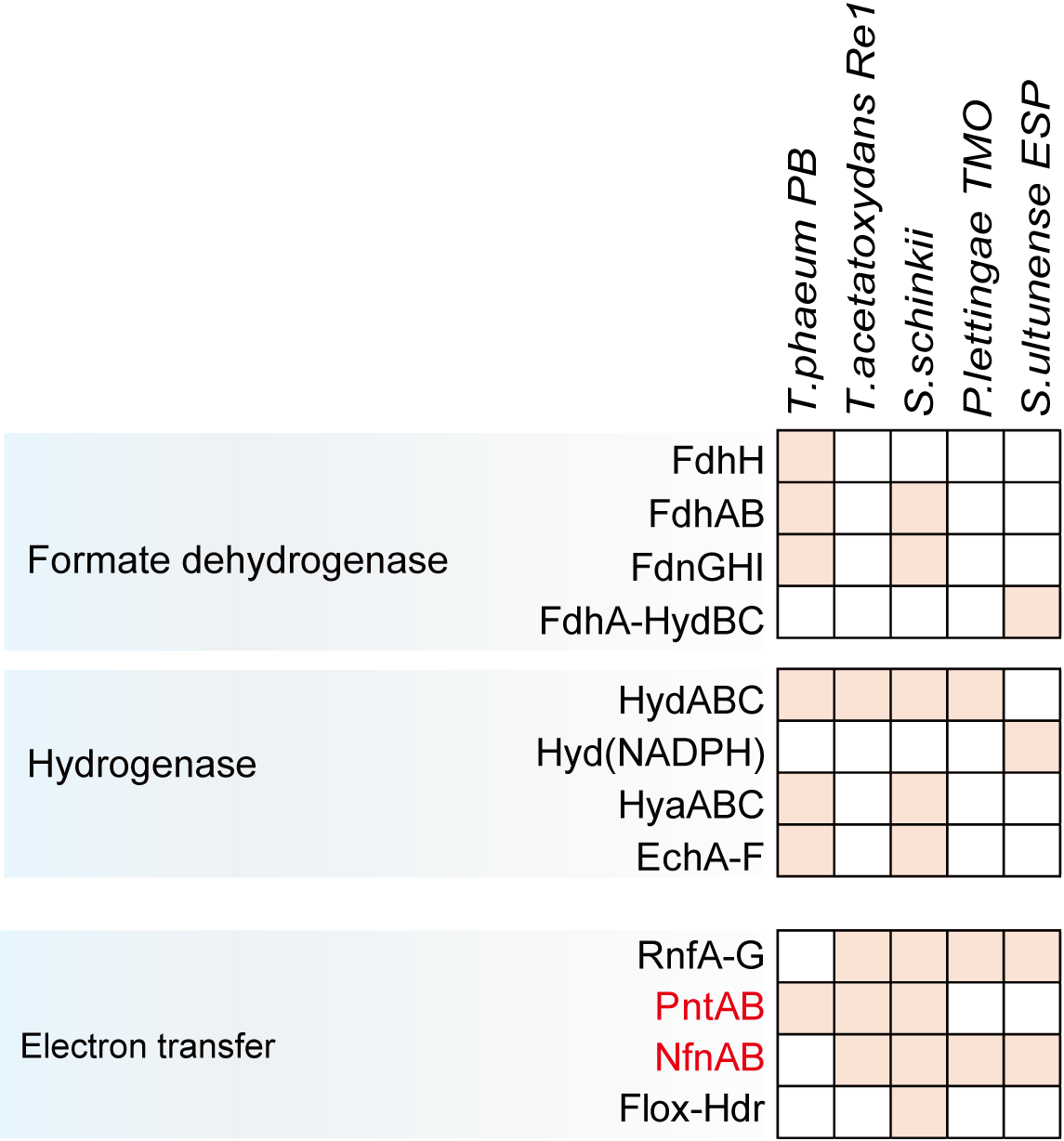

**Fig. S8.** Energy conservation strategies of known SAOB. Enzyme abbreviations and their corresponding genes are elaborated in Supporting Information Tables S7.

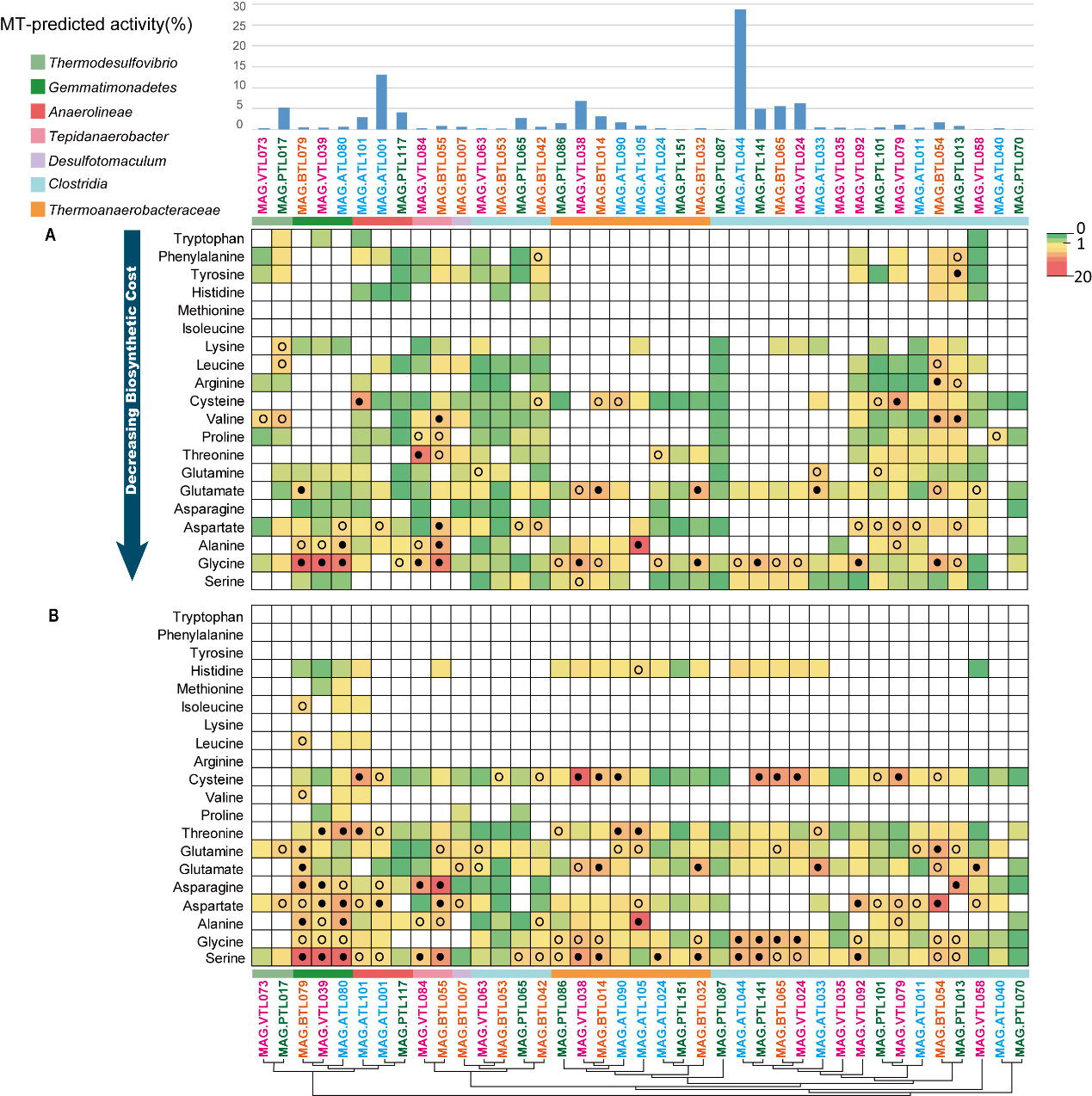

**Fig. S9.** Gene expression levels for amino acid biosynthesis (A) and degradation (B) of syntrophs in thermophilic chemostats. For each MAG, the percentages of the metatranscriptomic (MT) reads mapped to the MAG out of the metatranscriptomics mapped to all MAGs (both *Bacteria* and *Archaea*) are shown. The gene expression levels are calculated as reads per kilobase of transcript per million reads mapped to individual MAG (RPKM) normalized to the median gene expression for the corresponding MAG (RPKM-NM) averaged from duplicate samples. Pathways containing genes with RPKM-NM greater than the octile and quartile are marked (filled and open dots, respectively). Enzyme abbreviations and their corresponding genes are elaborated in Supporting Information Table S8-S9.

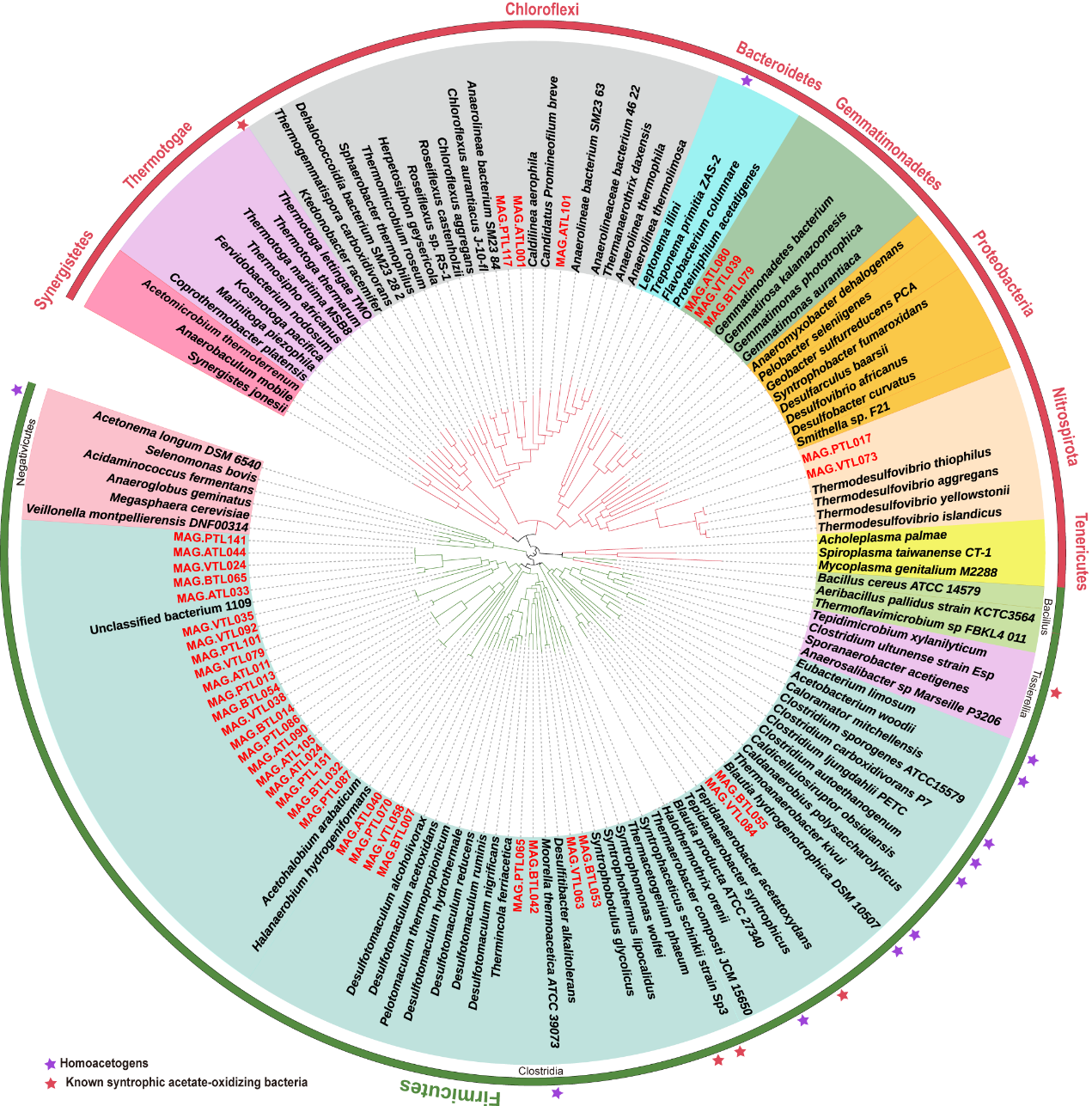

**Fig. S10.** Phylogenetic analyses of MAGs of syntrophs in thermophilic chemostats.

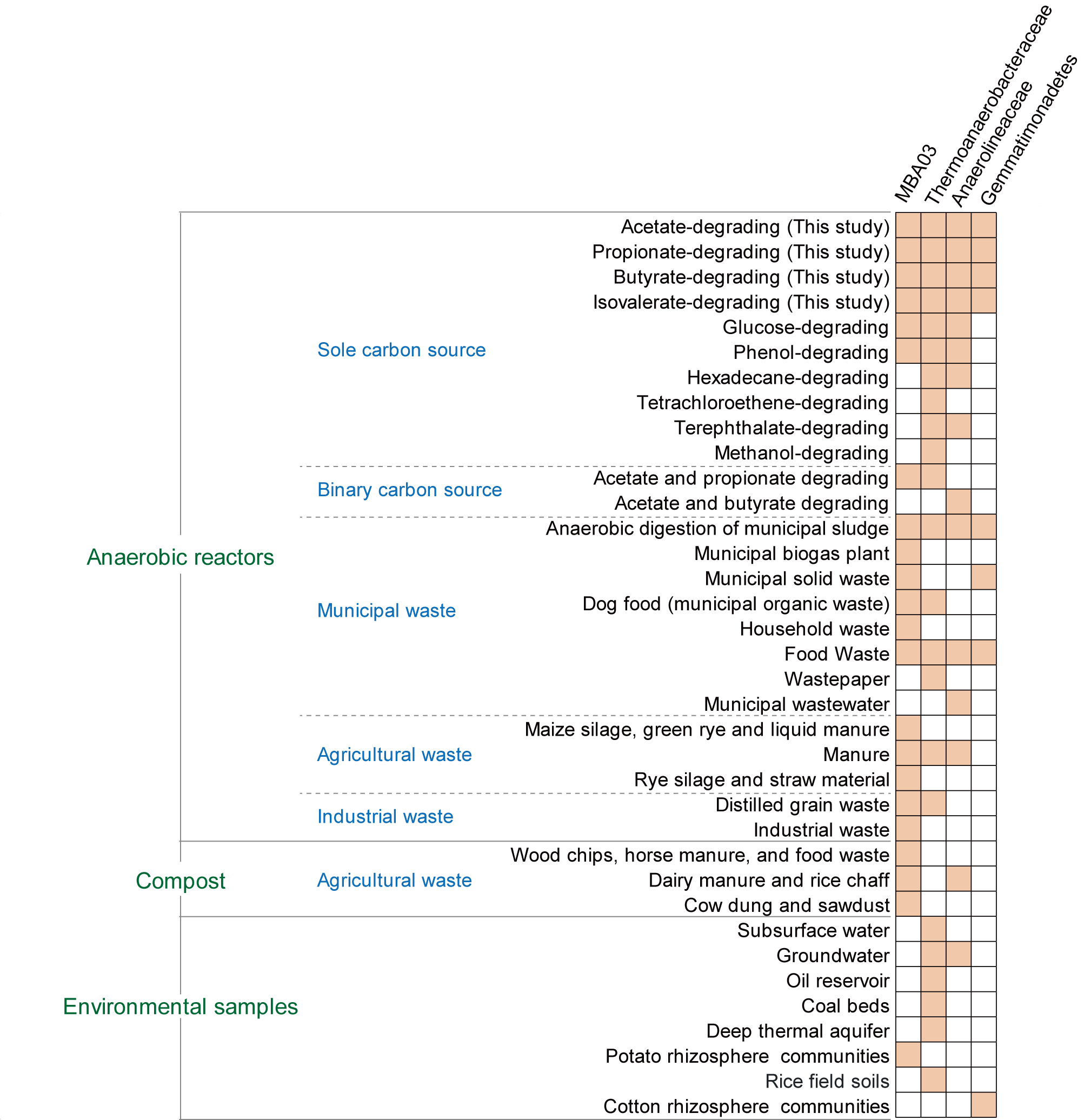

**Fig. S11.** Ecological characterization of the novel acetate oxidizer. The orange tiles represent environments from which sequences related closely (≥ 97% similarity) to the novel acetate oxidizer in this study.

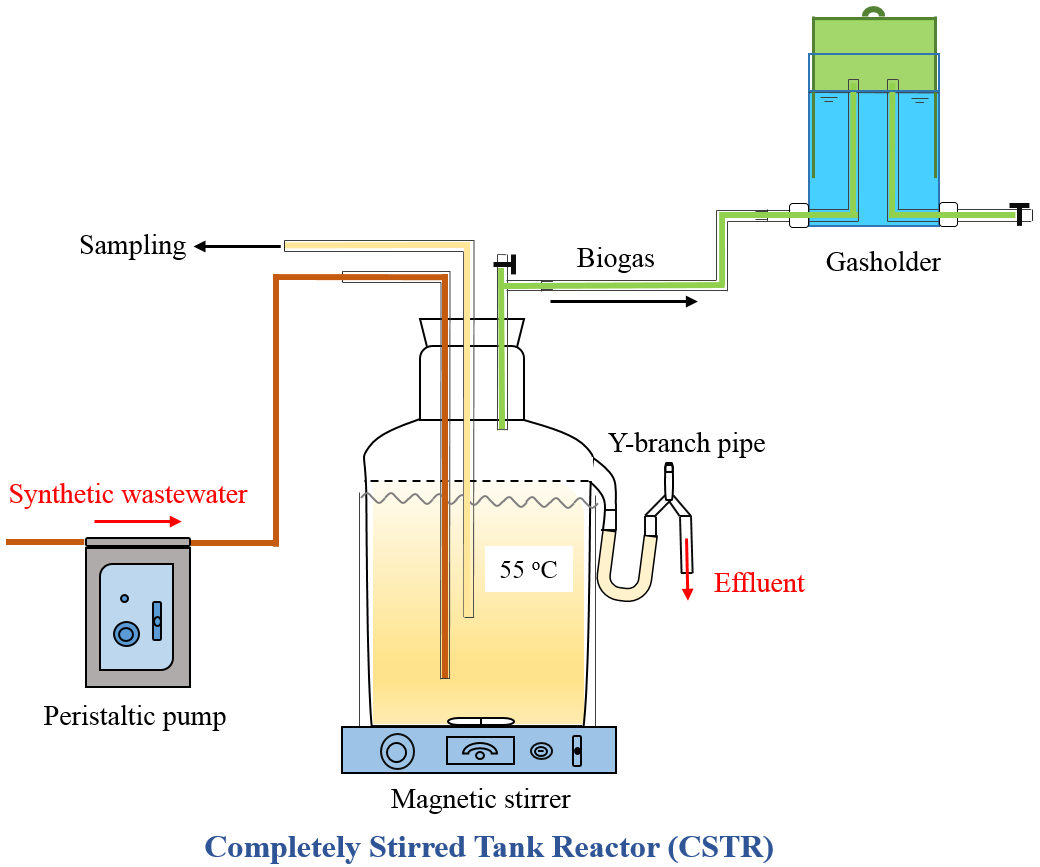

1.8L

**Fig. S12.** Schematic diagram of the thermophilic completely stirred tank reactor.
